# Genome-wide molecular recording using Live-seq

**DOI:** 10.1101/2021.03.24.436752

**Authors:** Wanze Chen, Orane Guillaume-Gentil, Riccardo Dainese, Pernille Yde Rainer, Magda Zachara, Christoph G. Gäbelein, Julia A. Vorholt, Bart Deplancke

## Abstract

Single-cell transcriptomics (scRNA-seq) has greatly advanced our ability to characterize cellular heterogeneity in health and disease. However, scRNA-seq requires lysing cells, which makes it impossible to link the individual cells to downstream molecular and phenotypic states. Here, we established Live-seq, an approach for single-cell transcriptome profiling that preserves cell viability during RNA extraction using fluidic force microscopy. Based on cell division, functional responses and whole-cell transcriptome read-outs, we show that Live-seq does not induce major cellular perturbations and therefore can function as a transcriptomic recorder. We demonstrate this recording capacity by preregistering the transcriptomes of individual macrophage-like RAW 264.7 cells that were subsequently subjected to time-lapse imaging after lipopolysaccharide (LPS) exposure. This enabled the unsupervised, genome-wide ranking of genes based on their ability to impact macrophage LPS response heterogeneity, revealing basal *NFKBIA* expression level and cell cycle state as major phenotypic determinants. Furthermore, we show that Live-seq can be used to sequentially profile the transcriptomes of individual macrophages before and after stimulation with LPS, thus enabling the direct mapping of a cell’s trajectory. Live-seq can address a broad range of biological questions by transforming scRNA-seq from an end-point to a temporal analysis approach.

## Introduction

Most biological processes are inherently transient and dynamic, with cell states changing according to internal programs and/or external stimuli, as illustrated, for example, by the fact that the many different cell types in our body are derived from a single zygote. It is thus critical to not only understand a cell’s current state, but also how a cell arrived at that state, i.e. its molecular history, which ensures normal development or causes disease when deviating from its canonical path^1^. However, despite its importance, revealing the history of a cell remains an outstanding technological challenge. Several approaches have been developed with the intent of exploring a cell’s past, such as engineered genetic circuits^2^, recombinase- and CRISPR-based DNA editors^3–7^ and live cell imaging^8, 9^. While these methods represent exciting advances, they are limited by their ability to record only one or a few events per single cell rather than providing a comprehensive view of the molecular state of the cell^10^.

Single cell transcriptomics (scRNA-seq) is recognized as highly complementary to the above mentioned strategies, as it allows inferring the history and trajectory of cells based on unbiased transcriptomic read-outs^11–19^. Such scRNA-seq-based trajectory inference methods are mainly based on the principle that a snapshot of a population of cells at different stages can reflect the molecular changes of an individual cell over time. Although they have already provided numerous biological insights, these trajectory inference tools have a fundamental limitation in that there may be multiple potential dynamics for a measured distribution of cell states that give rise to them^20–22^. Thus, the models generated by these tools should be interpreted as statistical expectations rather than the actual transition path of the cells^23^. This in part explains why distinct cellular trajectory approaches often predict different trajectories on the same dataset^24^, and rationalizes why accurate trajectory inference remains one of the recently defined grand challenges in single-cell data science^25^. Moreover, while scRNA-seq technologies have undergone a remarkable transformation, from the initial profiling of only a few cells^26^ to processing thousands of cells or more at a time today^27, 28^, these technologies all share the same experimental drawback: they only allow end-point measurements since they require cell lysis and are thus destructive.

Here, we introduce Live-seq, which enables transcriptome profiling of living cells after cytoplasmic biopsy. The approach is based on fluidic force microscopy (FluidFM)^29, 30^, which we optimized to collect cytoplasmic mRNA and coupled to an in-house devised, highly sensitive low-input RNA-seq strategy. We demonstrate that Live-seq allows the profiling of single cell transcriptomes while keeping cells alive, thus enabling downstream functional analyses directly on the assayed cells. Consequently, Live-seq can function as a transcriptomic recorder, linking the “past” transcriptome of a cell to its downstream cellular phenotype. In addition, we demonstrate that Live-seq can be used to sequentially profile the transcriptomes of the same individual cells, providing a direct read-out of cellular dynamics. To broadcast the power of Live-seq, we recorded the transcriptomes of individual macrophage-like RAW 264.7 cells to identify the principal factors underlying macrophage LPS response heterogeneity, uncovering basal NFKBIA expression level and cell cycle state as major phenotypic determinants.

## Results

### The basics of Live-seq

The total RNA content in mammalian cells is estimated to be ∼10 pg, ranging from as little as 1 pg up to 50 pg, dependent on cell type^31^. Because FluidFM can be used to sample a fraction of the total RNA content of a cell in a tunable way, we reasoned that cytoplasmic biopsies that preserve cell viability would yield RNA at a scale of a few picograms down even to sub-picograms. Our first challenge was therefore to optimize the FluidFM RNA extraction procedure with the goal of making transcriptome profiling compatible with such small amounts of RNA input. To do so, we first set out to minimize the degradation of sampled RNA and the loss of (already picoliter-scale) sample during the cell-to-tube transfer by i) reducing the extraction time, ii) lowering the temperature, iii) implementing a preloading of the FluidFM probe with sampling buffer with the goal of immediately mixing the extracted cytoplasmic fluid with RNase inhibitors, iv) releasing the extract into a microliter droplet containing buffer that is compatible with downstream RNA-seq, v) introducing a washing step to avoid cross-contamination, and vi) implementing image-based cell tracking for sequential extraction (relevant for sequential Live-seq as discussed below). The process is detailed in **Figure 1a** and **Methods**.

**Figure 1.**
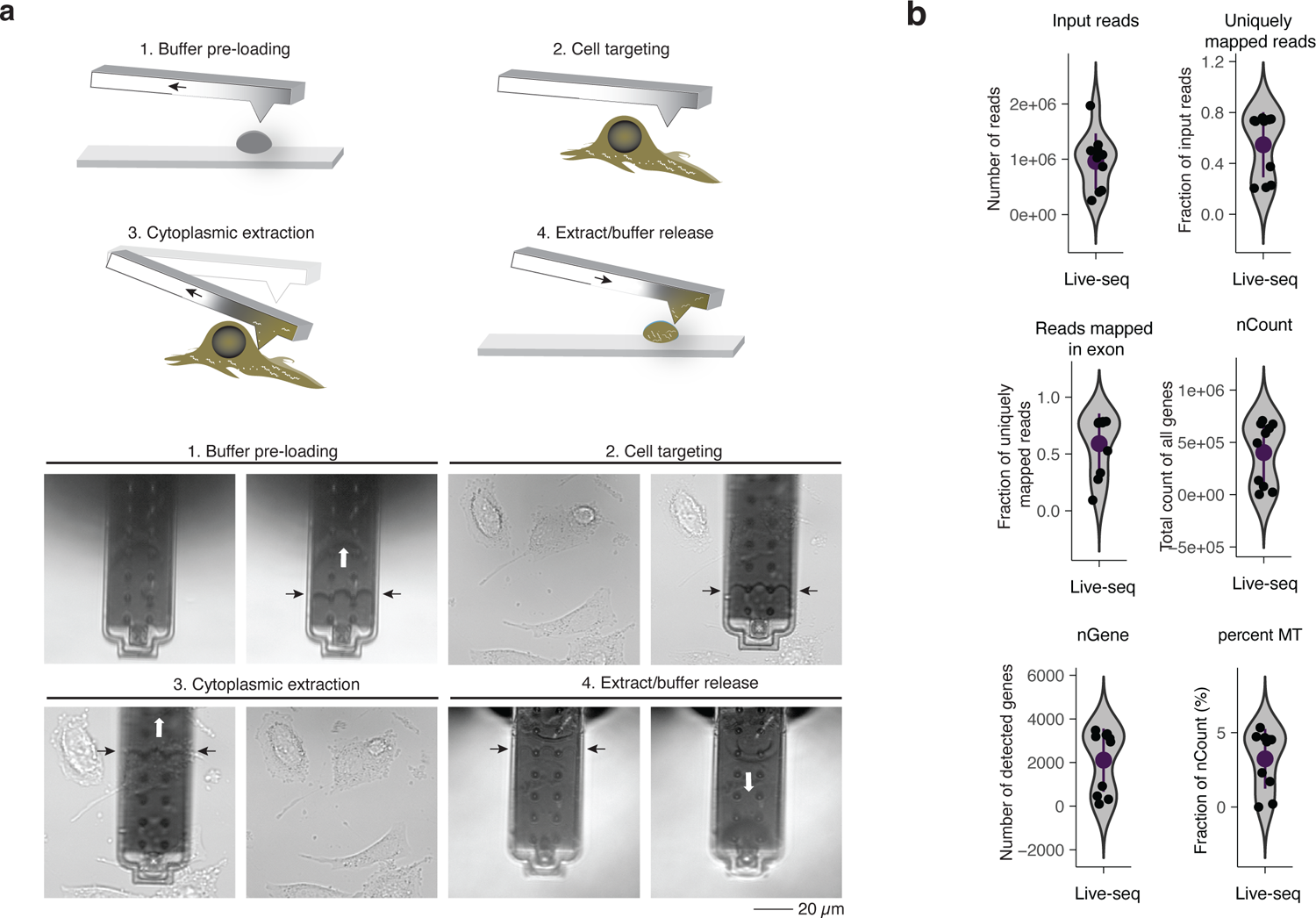
Live-seq combines optimized FluidFM-based live cell biopsy with enhanced Smart-seq2 RNA-seq. (**a**) The upper panels illustrate the Live-seq sampling procedure using FluidFM and the lower panels are the representative images (here, applied on pre-adipocyte IBA cells). The white arrows indicate the application of under-/over-pressure. The black arrows indicate the level of buffer and extract in the probe. (**b**) Quality control of Live-seq applied on IBA cells based on the parameters that are listed above each panel. N = 10 cells. nGene: number of detected genes. nCount: total count of all genes. Percent MT: percentage of counts from mitochondrial genes. Error bars represent mean +/- SD.

Since Smart-seq2^32^ was widely appreciated as one of the most sensitive RNA-seq methods to detect low amounts of RNA^28^ at the time of method development, we assessed whether it could amplify cDNA at the anticipated picogram scale. While successful for RNA inputs above 5 pg, little to no cDNA was recovered from inputs of 2 pg or less (**Supplementary Figure 1a**). To improve recovery, we systematically assessed and subsequently optimized the efficiency of each step in the workflow (detailed in **Supplementary Text 1**). These efforts yielded a protocol that enables the reliable detection of 1 pg of total RNA by selecting a dedicated reverse transcription enzyme (Maxima H Minus Reverse Transcriptase), limiting adaptor concatemer formation by incorporating 5’ biotinylated template-switching^33, 34^ and oligo-dT oligonucleotides, and optimizing the concentration of these components (**Supplementary Figure 1a-g, Methods**). Readouts of several parameters attest to the high sensitivity of the method (**Supplementary Figure 1g-i**)^32^, including i) the high rate of uniquely mapped reads, ii) the number of detected genes, and iii) the gradual increase of the cumulative proportion of each library assigned to the top-expressed genes from 1 pg input RNA, while absent in the negative control (0 pg input RNA). Consistent with this, we observed that the reads from the negative control are overrepresented by poly A and TSO sequence stretches and that they map to only few genes (**Supplementary Figure 1j-k**).

Next, we combined FluidFM-based cytoplasmic sampling with our enhanced, low-input RNA profiling workflow to probe transcriptomes from living cells. As an initial proof of concept, we sampled mouse immortalized brown preadipocyte (IBA) cells^35^ and detected on average about 2100 genes (nGene) at a sequencing depth of around 1 million reads per sample (**Figure 1b**). Since the same FluidFM probe can be used for sequential sampling of multiple cells, a wash process was implemented to prevent cross-contamination (>99% accuracy based on read mapping) (**Supplementary Figure 1l, Methods**).

Together, these technological advances constitute the methodological foundation of Live-seq, which has demonstrated potential for transcriptome profiling using cytoplasmic biopsies.

### Live-seq enables the stratification of cell types and states

To assess the cell identity- or even cell state-resolving power of Live-seq-derived cell transcriptomes, we applied our approach to distinct cell types. These included IBA cells, primary mouse adipose stem and progenitor cells (ASPCs), as well as two monocyte/macrophage-like RAW264.7 cell lines: one wildtype and one RAW264.7 subline (RAW-G9) containing an mCherry reporter under the control of the *Tnf* promoter^36^ to facilitate downstream functional analyses (shown below). The latter cell line and its parental RAW264.7 line are molecularly and phenotypically similar^36^ and will therefore be further referred to as “RAW” cells. To investigate the capacity of Live-seq to study cell state transitions, RAW cells were treated with lipopolysaccharides (LPS), an outer membrane component of Gram-negative bacteria capable of activating monocytes and macrophages (**Figure 2a**), or mock-treated as a control. In total, we harvested 443 samples across four replicates and generated 390 libraries for sequencing (**Supplementary Figure 2a** and **Methods** for a description of the different datasets). From these, 175 samples passed our filtering criteria, featuring i) more than 1000 genes/sample, ii) less than 30% of reads derived from mitochondrial genes, and iii) more than 30% uniquely mapped reads (**Supplementary Figure 2a** and **Methods**). Similar to Smart-seq2^32^, reads spanned the full length of transcripts, with the 5’ and 3’ ends being less covered (**Supplementary Figure 2b**). An average of 3,218 genes were detected with further quality controls shown in **Supplementary Figure 2c-e**.

**Figure 2.**
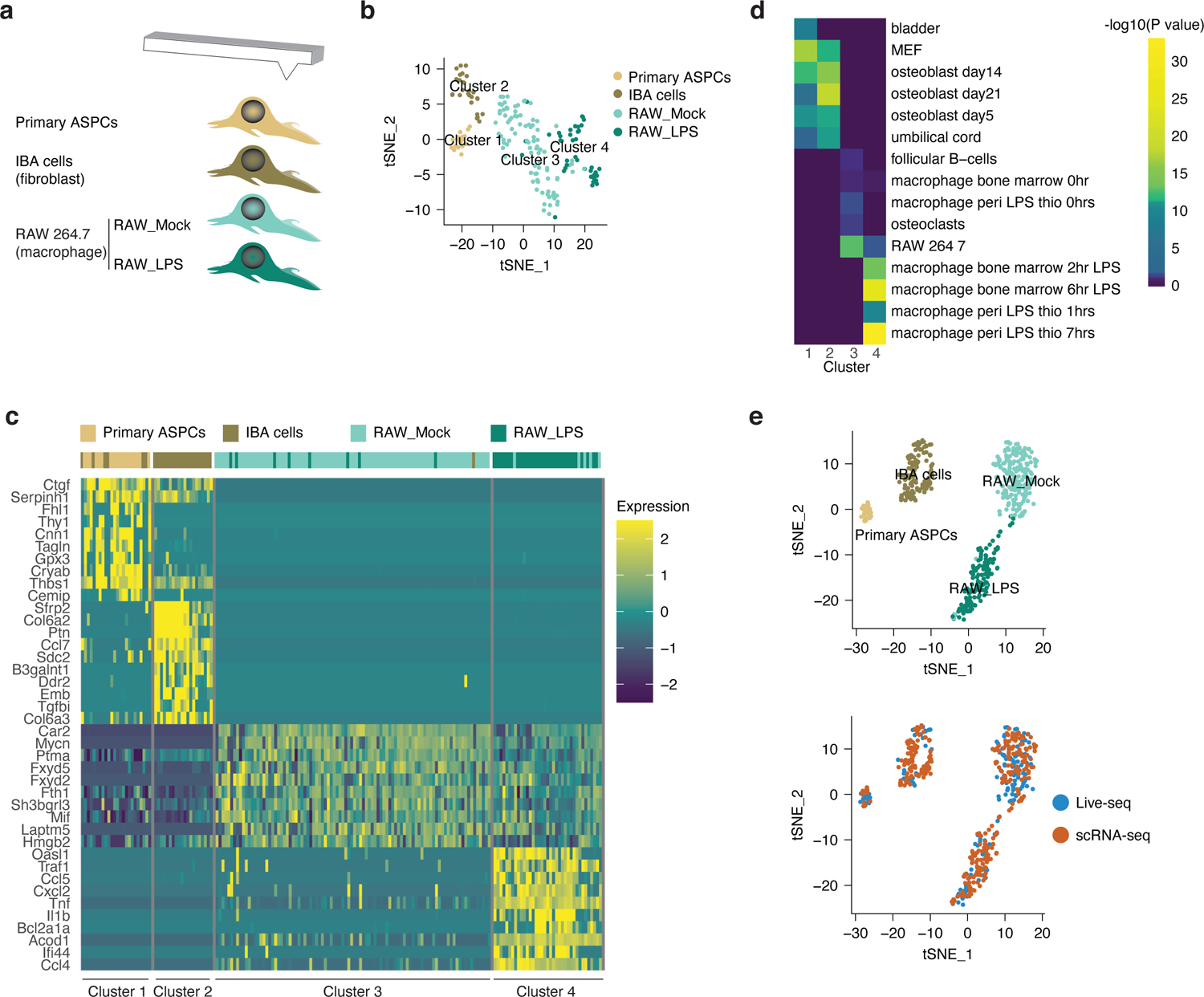
Live-seq enables the stratification of cell type and treatment (cell state). (**a**) Experimental setup. 100 nM LPS or PBS was used for RAW_LPS and RAW_Mock, respectively. (**b**) Clustering of Live-seq samples based on the top 500 highly variable genes as visualized in a tSNE plot. (**c**) Heatmap showing the top 10 differentially expressed genes, stratified according to cell type or treatment (state). (**d**) Mouse gene atlas-based prediction of cell type/state of each cluster using the top 100 marker genes. (**e**) Visualization of Live-seq and scRNA-seq data after anchor-based data integration (**Methods**) reveals no obvious molecular differences. tSNE projection of the integrated Live-seq and scRNA-seq data according to cell type and treatment (upper panel) and approach (lower panel).

We observed that our Live-seq data separates into four clusters (**Methods**). Cell type and treatment thereby largely dominated over library complexity (number of genes detected), replicate identity, or RAW cell subline (**Figure 2b** and **Supplementary Figure 2f**), supporting Live-seq’s ability to effectively profile both primary and cultured cells. Cluster 1 mainly represented transcriptomes from primary ASPCs, Cluster 2 from IBA cells, Cluster 3 from RAW cells without LPS treatment, and Cluster 4 from RAW cells exposed to LPS. Hierarchical clustering revealed consistent results (**Supplementary Figure 2g**), implying clustering robustness.

To validate the biological relevance of these four clusters, we performed differential gene expression analysis (**Supplementary Table 1**), yielding results that were largely consistent with the cellular characteristics of each cluster (**Figure 2c**, **Methods**), i.e. mesenchymal stem cell genes in Cluster 1, mesenchymal/fibroblast genes in Cluster 2, immune-related genes in Clusters 3 and 4 with the latter also featuring genes induced by immune stimulation. Furthermore, Gene Ontology (GO) term enrichment analysis revealed that terms related to extracellular matrix organization and extracellular matrix disassembly are enriched in Clusters 1 and 2 (**Supplementary Figure 2h**), whereas Cluster 3 showed enrichment for antigen receptor−mediated signaling pathway- and leukocyte differentiation-related terms, and Cluster 4 for immune-related terms such as inflammatory response and response to LPS. In addition, matching of the top 100 marker genes to mouse gene atlas annotations^37^ confirmed the expected molecular similarity of Cluster 1 and 2 cells to mouse embryonic fibroblasts (MEF) and osteoblasts (**Figure 2d**), reflecting their mesenchymal /fibroblast nature. Cluster 3 cells were correctly annotated as RAW264.7/macrophages without treatment, and Cluster 4 cells as macrophages with LPS treatment (**Figure 2d**).

Finally, to evaluate whether cytoplasmic mRNA biopsies can act as suitable representations of full (lysed) cell transcriptomes, we compared Live-seq gene expression profiles to those obtained using the whole cell Smart-seq2 assay (**Methods**). Using the same cell types and treatment groups, we sequenced 343 cells with scRNA-seq, with 338 cells passing the same filtering criteria applied to Live-seq samples (**Supplementary Figure 2i**). We detected, as expected, a greater average of genes per cell compared to Live-seq: 7,952 genes at a sequencing depth of around 700K reads per cell with the quality control parameters attesting to high data quality (**Supplementary Figure 2j-k**). Graph-based clustering partitioned these cells also into four groups (**Methods**), accurately capturing the underlying cell type and/or treatment group (**Supplementary Figure 2l**). Results obtained by differential gene expression analysis (**Supplementary Figure 2m** and **Supplementary Table 2**) followed by GO analysis (**Supplementary Figure 2n**) were in line with those obtained by Live-seq (**Figure 2d**) in that the four clusters correctly annotated mesenchymal/fibroblast-like cells and mock-treated or LPS-treated macrophages, respectively (**Supplementary Figure 2o**). We then integrated the Live-seq and scRNA-seq data to explore putative molecular differences between the two conceptually distinct approaches. Even without batch correction, we already observed a dominance of cell type/state signals over those stemming from the sampling approach (**Supplementary Figure 2p**). Correlation analysis further revealed that a meta read-out of cells (i.e. collapsing all cells into a virtual bulk sample) produced the expected pattern with the Live-seq ASPC gene expression profile correlating highest with the scRNA-seq-based one and the same for IBA cells, RAW cells with and without LPS treatment, respectively (**Supplementary Figure 2q**). We finally applied a single-cell-tailored canonical correlation analysis (CCA) and mutual nearest neighbors (MNNs)-based approach that is embedded in Seurat^38^ to remove any potential technical batch effects. This further consolidated the clustering of cells according to cell type and state regardless of the sampling approach (**Figure 2e** and **Supplementary Figure 2r, Methods**) and sequencing depth (**Supplementary Figure 2s**).

Together, these results demonstrate that Live-seq enables the accurate stratification of cell types and states similar to conventional scRNA-seq, but without the need to lyse cells.

### Live-seq preserves cell viability, transcriptome, and growth

The analyses described above allowed us to benchmark Live-seq to a widely used conventional, whole cell transcriptome profiling approach (**Methods**). However, Live-seq was designed to be minimally invasive and to conserve cell viability to allow for continuous monitoring of the very same cell. To address this capacity, we first evaluated cell viability after sampling (**Methods**). While the different cell types that were sampled exhibited considerable cell size differences, the post-extraction viability was similar for all the cell types after Live-seq sampling. Indeed, overall cell viability ranged between 85 and 89% with an average extraction volume of 1.1 pL (**Supplementary Figure 3a** and **Methods**) compared to a viability of 90-95% after conventional trypsin-based cell dissociation (data not shown).

These high cell viability numbers suggest that cells are able to quickly recover after a portion of their cytoplasm has been removed. However, assessing cell viability does not inform on putative, short-term molecular effects that Live-seq may impose on targeted cells. To address this, we used standard scRNA-seq (**Methods**) to profile the individual transcriptomes of IBA cells 1 hour and 4 hours post-cytoplasmic sampling (**Figure 3a**), while using non-probed cells as negative controls. Quality controls on the cDNA and sequencing libraries (such as mapping rate, number of detected genes) were indistinguishable among the three cell groups (control, 1h and 4h post-Live-seq-sampling; **Supplementary Figure 3b-c**). Remarkably, data analysis revealed no distinct condition-related clusters (**Figure 3b**), further supported by the observation that only 12 genes were found to be significantly differentially expressed (DE) among the three conditions (**Figure 3c**). To assess subtle changes in biological processes that may not be visible at the DE level (because of hard cut-offs), we performed a Gene Set Enrichment Analysis (GSEA) on GO biological process terms using the expression levels of all genes, but this did not reveal any specifically enriched processes.

**Figure 3.**
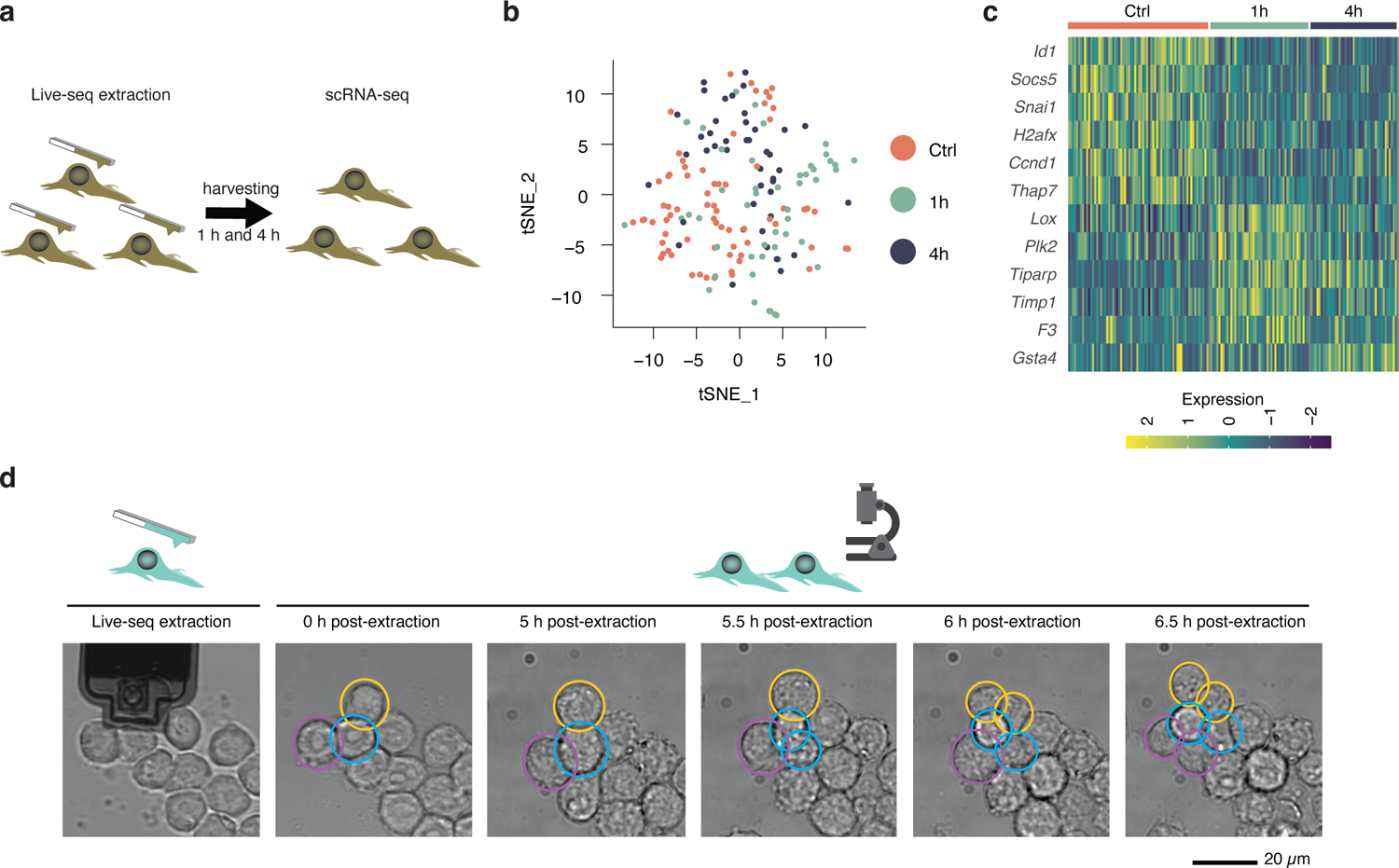
Live-seq preserves a cell’s viability, transcriptome, and growth. (**a**) Schematic of the experimental design to evaluate putative transcriptomic changes after Live-seq extraction. IBA cells were first extracted after which they were harvested and subjected to scRNA-seq 1 hour and 4 hours post-extraction. Cells that were not extracted were included as control. (**b**) A tSNE projection of control cell scRNA-seq data (Ctrl, 70 cells) as well as 1 hour (49 cells) and 4 hours (43 cells) post Live-seq extraction scRNA-seq data does not reveal clearly distinct clusters based on the top 500 most variable genes. (**c**) All of the genes (12) that were found differentially expressed between the control and Live-seq-sampled cells. (**d**) Representative time-lapse images showing the division of RAW cells post-Live-seq extraction. Three dividing cells are outlined in colored circles. The cell in the yellow circle was subjected to cytoplasmic extraction, while those in purple and blue were not.

These experiments indicate that Live-seq does not induce major, short-term gene expression alterations. To assess putative longer-term phenotypic effects and thus to study the recovery of Live-seq-probed cells in greater detail, we examined the growth dynamics of RAW cells after Live-seq using time-lapse microscopy. We focused on RAW cells for two reasons: i) their size is smallest among the three sampled cell types, and thus their behavior may be most impacted by a cytoplasmic extraction; ii) as semi-adherent cells, their almost spherical shape allowed for a straightforward approximation of cell volume changes over time. Using longitudinal cell volume monitoring, we found that extracted RAW cells are able to recover their volume quickly and continue to grow (**Supplementary Figure 3d-e**). Moreover, while overall cell behavior was variable, we observed that some cells divided already a few hours post-Live-seq sampling, in similar fashion to non-extracted cells (**Figure 3d** and **Supplementary Figure 3d-e**). Although we cannot completely rule out that Live-seq introduces a small cell cycle delay, as cell size and cell division tend to be intrinsically coupled^39^, our results indicate that cells still progress through the cell cycle even after having been subjected to a cytoplasmic biopsy.

Altogether, we conclude that Live-seq enables the profiling of cell transcriptomes without imposing major perturbations on a cell’s basic properties such as viability, transcriptome, or growth, thus opening a new avenue to link a cell’s molecular state directly to its present and future phenotypic properties.

### Live-seq records molecular events that are predictive of a cell’s downstream phenotype

It is well recognized that macrophages, including the RAW cells already used above, respond to LPS in a heterogeneous fashion^36^. Although this has been the subject of intense interrogation^40^, a systematic, genome-wide analysis of what molecular factors drive this heterogeneity has not been performed. To do so, we used Live-seq to profile the transcriptomes of individual RAW cells to link the molecular state of each macrophage to its downstream, LPS-induced phenotype. Specifically, we first examined single RAW cells with Live-seq to record their respective transcriptomes in the ground-state. We then subjected the same cells to LPS treatment and tracked LPS-induced *Tnf* promoter-driven mCherry expression (hereafter referred to as *Tnf*-mCherry) using live cell-imaging (**Figure 4a**). We observed that *Tnf*-mCherry intensity increased upon LPS treatment, although this intensity varied greatly between RAW cells (**Figure 4b**), consistent with previous findings^36^. Importantly, no significant difference in the heterogeneity of *Tnf*-mCherry intensity profiles was observed between cells subjected to Live-seq sampling and those that were not (**Figure 4b**, two-sided Wilcoxon rank-sum test, *P* = 0.876). These results provide additional support to the notion that Live-seq does not markedly affect a cell’s phenotype (**Figure 3**).

**Figure 4.**
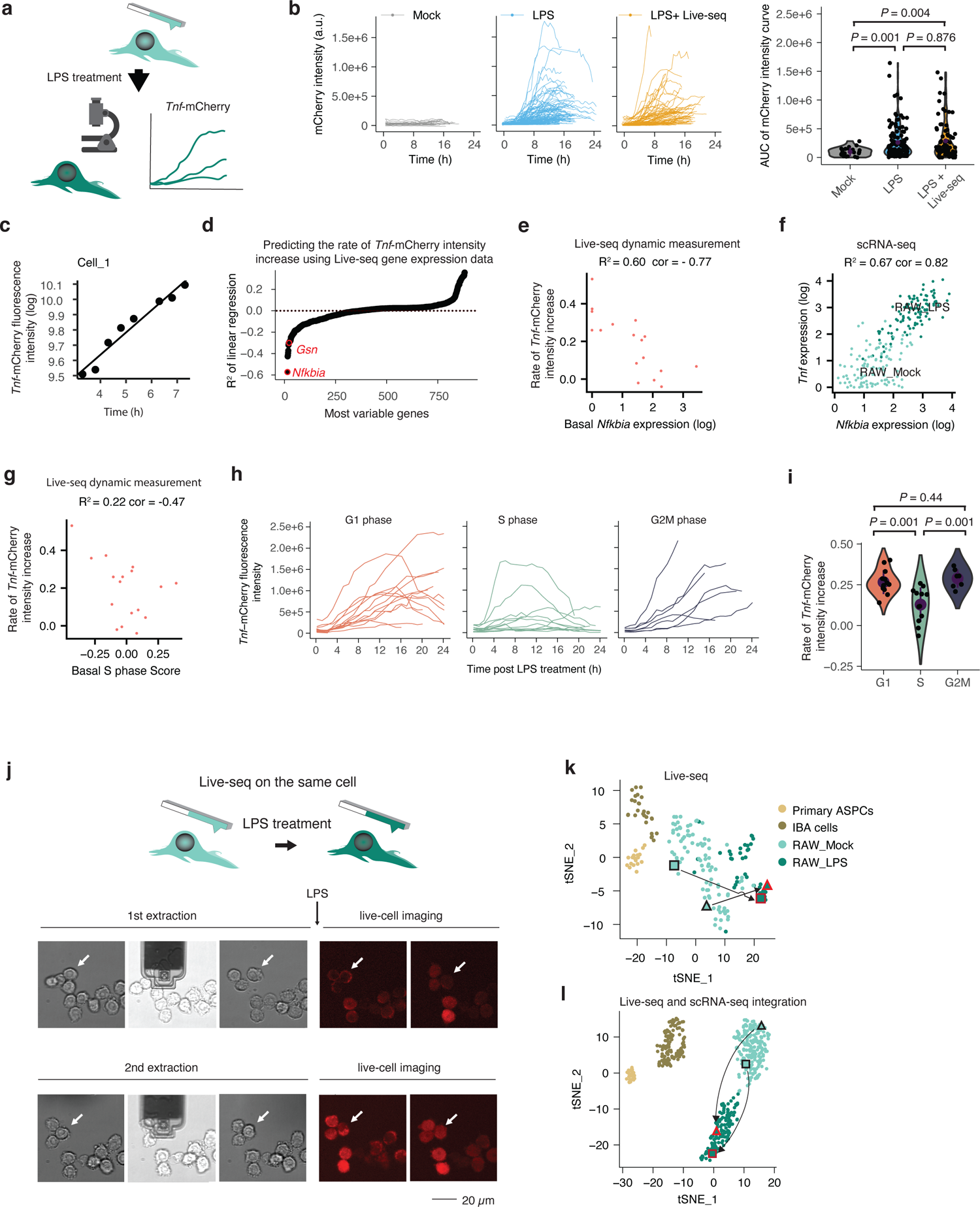
Live-seq enables longitudinal single-cell analyses. (**a**) Schematic illustrating the coupling of Live-seq with live-cell imaging. RAW cells were first subjected to Live-seq. Subsequently, they were treated with LPS after which *Tnf*-mCherry fluorescence was tracked by time-lapse imaging. (**b**) (Left three panels) mCherry intensity profiles of LPS-treated control cells or those subjected to Live-seq sampling. Mock-treated cells were used as negative control. (Right panel) Area Under the Curve (AUC) values for all cells shown in the left panels were plotted according to the indicated conditions. N = 23 cells for the Mock condition, 122 cells for the LPS condition and 77 cells for the LPS + Live-seq condition. (**c**) The time post-LPS treatment (within a window of 3 to 7.5 hours) and the log transformed *Tnf*-mCherry fluorescence intensity shows a linear relationship in individual cells (one representative cell is shown here) with the intercept representing the *Tnf*-mCherry basal level and the slope the rate of fluorescence intensity increase. Please see **Supplementary** Figure 4a for similar data on all other cells. (**d**) A linear regression model was used to predict the slope (the rate of *Tnf*-mCherry fluorescence intensity increase) calculated from data shown in (c) and **Supplementary** Figure 4a based on ground-state gene expression data recorded by Live-seq. The most variable genes from both Live-seq and scRNA-seq data were used, while genes that were not detected in Live-seq samples were removed. Two genes, *Nfkbia* and *Gsn*, known to inhibit LPS-induced *Tnf* expression, are highlighted. (**e**) Basal *Nfkbia* expression, as determined by Live-seq, anti-correlates with the rate of *Tnf*-mCherry fluorescence intensity increase. R^2^ and Pearson’s r values are listed. (**f**) The expression of *Nfkbia* and *Tnf* is highly correlated in conventional scRNA-seq data. R^2^ and Pearson’s r values are listed. (**g**) The (basal) cell cycle S phase score of Live-seq samples (inferred from respective transcriptomes) anti-correlates with the rate of *Tnf*-mCherry fluorescence intensity increase, suggesting that cells in S phase respond weaker to LPS treatment. R^2^ and Pearson’s r values are listed. (**h**) *Tnf*-mCherry fluorescence intensity profiles in individual LPS-treated cells that are stratified here according to their respective cell cycle phase, as inferred from the Fucci cell cycle indicator miRFP709-hCdt1 that was transduced in RAW cells after which miRFP709 fluorescence was tracked for 24 hours prior to LPS treatment (**Supplementary** Figure 4f, **Methods**). (**i**) The rate of *Tnf*-mCherry fluorescence intensity increase (slope) between 3 to 7.5 hours post-LPS treatment was calculated based on data shown in (h). G1 phase: 13 cells; S phase: 12 cells; G2M phase: 7 cells. (**j**) Sequential Live-seq sampling on the same cells. Schematic illustrating the sequential sampling procedure, and time-lapse bright-field and mCherry fluorescence images of sequential Live-seq sampling (i.e. sequential extractions). The arrow points to the sequentially sampled cell. (**k**) tSNE-based visualization of Live-seq data highlighting the two sequentially sampled cells by a triangle and rectangle, respectively. A black shape outline represents unstimulated cells, whereas a red shape outline represents LPS-stimulated cells. (**l**) tSNE-based visualization of integrated scRNA-seq and Live-seq data, highlighting the transition of two sequentially sampled cells from one state to another. The annotation of these cells is as described in (k). For (b,i), error bars represent the mean +/- SD and *P* values were determined by a two-sided Wilcoxon rank-sum test.

Given the canonical heterogeneous response of RAW cells to LPS (**Figure 4b**), we then set out to uncover the principal ground-state molecular factors (as derived by Live-seq) that drive this heterogeneity. To do so, we used a linear regression model correlating the expression of Live-seq-detected genes within individual cells with the corresponding *Tnf*-mCherry response profiles. Specifically, given that we observed that the log form of each response profile is a linear function during a 3 to 7.5 hour post-LPS treatment time window (**Figure 4c** and **Supplementary Figure 4a**), we aimed to use ground-state gene expression data to predict the two principal *Tnf*-mCherry profile parameters: the basal *Tnf*-mCherry expression (intercept) and the rate of fluorescence intensity increase (slope) (**Figure 4d** and **Supplementary Figure 4b**). Applying the linear model revealed *Tnf* as the second-best predictor (R^2^ = 0.50) of basal *Tnf*-mCherry intensity (intercept), validating the accuracy of Live-seq, with an uncharacterized long non-coding RNA *4632427E13Rik* as the best predictor (**Supplementary Figure 4b** and **Supplementary Table 3**). Next, we set out to identify factors that predict both the dynamics and amplitude to which a cell responds to LPS as defined by the rate of *Tnf*-mCherry intensity increase (slope). Using the Live-seq data, we generated an unsupervised, genome-wide rank of factors according to which extent they impacted a macrophage’s ability to respond to LPS. We thereby identified *Gelsolin* (*Gsn*) as one of the top negatively correlating factors (R^2^ = 0.33, Pearson’s r = −0.58) (**Supplementary Figure 4c**), consistent with its role in suppressing LPS-induced *Tnf* expression^41^, likely via direct LPS inactivation^42^. However, the strongest negative predictor of the rate of *Tnf*-mCherry intensity increase was *Nfkbia* (also termed NF-kappa-B inhibitor alpha, *IkBα*; R^2^ = 0.60, Pearson’s r = −0.77) (**Figure 4d-e** and **Supplementary Table 4**). This observation seems counterintuitive at first, given the positive correlation between *Nfkbia* and *Tnf* expression in conventional snapshot scRNA-seq data (**Figure 4f**). However, it is in fact consistent with the well-known function of NFKBIA as a key negative regulator of the LPS-NF-κB signaling pathway by suppressing NF-κB, which itself regulates the co-expression of both *Nfkbia* and *Tnf*^43^. These findings highlight the complementarity of the Live-seq-based dynamics approach and the scRNA-seq-based snapshot assay. Moreover, while many genes are known to regulate the LPS-NF-κB signaling pathway, including IKKs as positive regulators and A20 as a negative one^43^, the transcriptome-wide read-out enabled by Live-seq clearly points to *Nfkbia* expression as the strongest negative predictor of LPS-induced *Tnf* expression. Interestingly, we found that *Nfkbia* expression is also the most variable among the annotated NF-κB pathway components in multiple primary macrophage populations (**Supplementary Figure 4d**), further supporting the notion that NFKBIA acts as a principal driver of macrophage LPS response heterogeneity.

LPS has been shown to inhibit macrophage cell cycle progression^44^, which appears consistent with the observed anti-correlation between the conventional scRNA-seq-derived S phase score of RAW cells and *Tnf* expression (**Supplementary Figure 4e,** R^2^ = 0.46, Pearson’s r = −0.67). However, this anti-correlation may also reflect that a macrophage’s ability to respond to LPS is influenced by its cell cycle phase. To investigate these two plausible scenarios, we correlated the ground-state cell cycle phase of each cell, as inferred from pre-recorded Live-seq data, to its corresponding downstream LPS response profile. As shown in **Figure 4g**, this analysis revealed that cells tend to respond weaker to LPS when they are in S phase (R^2^ = 0.22, Pearson’s r = −0.47). To experimentally validate this observation, we expressed a fluorescent ubiquitination-based cell cycle indicator (Fucci) miRFP709-hCdt1^45^ in RAW cells (**Methods**). Prior to LPS stimulation, this indicator was tracked for 24 hours to determine the cell cycle phase of each cell (**Supplementary Figure 4f**), after which the LPS-induced *Tnf*-mCherry fluorescence intensity was measured. Since the Fucci system does not allow distinguishing the S from G2M phase, the boundary between the S and G2M phase was assigned by timing. As shown in **Figure 4h** and quantified in **Figure 4i** and **Supplementary Figure 4g**, cells in S phase responded significantly weaker (two-sided Wilcoxon rank-sum test, *P* =0.001 for both) to LPS stimulation (rate of of *Tnf*-mCherry intensity increase here) compared to their G1 and G2M phase counterparts, validating the Live-seq data. Together, these findings underscore the power of Live-seq to act as an unprecedented recording tool for the transcriptomes of individual cells and predict their phenotypic behavior.

### Sequential Live-seq cell sampling to measure cellular dynamics

A large number of computational approaches have been developed to infer cellular dynamics from one-time sampling data^24^. While clearly helpful, these approaches suffer from the fact that definition of the original molecular cell state is highly challenging in the absence of direct measurements^22^. To alleviate this issue, we sought to establish a proof-of-principle, sequential Live-seq sampling approach, which would allow us to record a cell’s molecular signature before and after cell state transitioning. To do so, we sampled RAW cells a first time, then stimulated them with LPS, after which we sampled the same cells a second time. In addition, the respective cells were also monitored by time-lapse microscopy after both sampling events (**Figure 4j**). Out of the 14 cells that were sequentially probed, we obtained transcriptome profiles that passed the quality control cutoff criteria for at least one of the sampling points for five cells, and for both of the two sampling points for two cells. While limited in number, these cells, each constituting two distinct points in the same t-SNE map (**Figure 4k**), provide to our knowledge the first direct, transcriptome-wide read-out of a cell’s trajectory. They revealed the transition of each cell from a pre-treatment (RAW_Mock) to a post-treatment (RAW_LPS) state. Moreover, given that the Live-seq and scRNA-seq data could be suitably integrated (**Figure 2e**), projecting the Live-seq sequential sampling information on top of the integrated data allowed us to unambiguously establish the correct trajectory of cells processed by conventional scRNA-seq (**Figure 4l**). This is in contrast to inference-based methods, which, based both on co-expression and RNA velocity, found several spurious connections between unrelated cell states on scRNA-seq data (**Supplementary Figure 4h-i**). Even a limited number of Live-seq sampled cells might thus provide a reference coordinate system to map cell trajectories of existing and future scRNA-seq data sets. It is thereby worth noting that the transcriptomes obtained from each sequential extraction were comparable (based on hierarchical clustering) to those of cells sampled only once (**Figure 4k-l** and **Supplementary Figure 4j**). This provides additional support to our observation that Live-seq does not impose major molecular perturbations (**Figure 3**). Taken together, sequential sampling with Live-seq allows the acquisition of transcriptomic dynamics from the same cell, thus providing a direct readout of cell state trajectories.

## Discussion

To date, scRNA-seq analyses have been endpoint analyses, which renders the resolution of cellular phenotypic variation or cell trajectories challenging^22^. To alleviate this, we established Live-seq by coupling an enhanced FluidFM-based live-cell biopsy technology to a highly sensitive RNA-seq approach. We showed that Live-seq is capable of distinguishing cell types and states, while keeping cells alive and functional based on the fact that cells proceeded in their cell cycle, remained LPS-responsive similar to control cells and exhibited only minor transcriptomic changes after cytoplasmic sampling. The search for new technologies that can improve our ability to infer cellular dynamics is not unique to this study^46^. This is for example illustrated by the recent development of i) methods that metabolically label newly synthesized RNA with the aim of recording transcriptional dynamics by differentiating old versus new RNA^11–13^ and ii) scRNA-seq-coupled lineage tracing, which superimposes clonal information on single cell transcriptomes to better resolve lineage relationships^47^. While constituting important advances, these approaches remain conceptually end-point methodologies. This makes downstream molecular or functional analyses on the same cell impossible and confounds studies aimed at better understanding cellular heterogeneity or cell dynamics^23^. The clear advantage of these recently developed approaches, however, is that they support high-throughput, while Live-seq will clearly benefit from further automation procedures that are currently under way. An increase in mRNA detection sensitivity may further increase Live-seq’s efficiency. A recent study that also aimed to improve the Smart-seq2 approach^48^ features several modifications that are in line with our enhanced Smart-seq2 method, including the use of Maxima H Minus Reverse Transcriptase and a biotin-blocked TSO oligo. A direct comparison with this approach was outside the scope of this study, but it is clear that any improvement in sensitivity of low input RNA-seq will benefit the further development of Live-seq.

Live-seq can be used to address a diverse range of biological questions, notably as a transcriptome-wide recording tool to resolve molecular determinants of cellular heterogeneity. Live-seq is in this regard complementary to other DNA-based recording systems that measure molecular events in a cell’s DNA after which these recordings can be recovered by DNA sequencing and subsequently linked to the downstream phenotype of that same cell^3–7^. The advantages of these DNA-based recording approaches include the stability of the recorded signal over time, the ability to sense biological signals *in cello*, and the fact that the recorded information can be read out in a large number of cells. However, disadvantages include the requirement to pre-engineer cells of interest, and difficulties in quantifying variability. Most importantly, current DNA-based recording systems register only one or a few events per cell^10^, while Live-seq provides a transcriptome-wide read-out. In this study, we used Live-seq to specifically study the molecular mechanisms underlying LPS response variation among macrophages (**Figure 4a-i**), which has been proposed to increase their response range to pathogens^40^. Multiple mechanisms such as stochastic fluctuations in gene induction^49^, differential cytoplasmic-nuclear distribution of transcription factors^50, 51^ and cellular environment^52^ have been proposed to contribute to this heterogeneity. However, a systematic assessment of whether and to which extent the molecular state of a cell impacts this heterogeneity had not yet been achieved. By exploiting Live-seq’s capacity to perform transcriptome-wide recording, we were able to generate a genome-wide, statistical ranking of a gene’s impact on influencing a macrophage’s response to LPS, revealing that the initial molecular state, and most prominently the expression levels of *Nfkbia* and *Gelsolin*, drive overall phenotypic heterogeneity (**Supplementary Table 4**). Moreover, by further mining the Live-seq data, we uncovered that the cell cycle phase also influences the LPS response, which we confirmed experimentally (**Figure 4g-i**). Transcription and DNA replication in a cell’s S phase are in inherent conflict, which can be a source of DNA damage and genome instability^53^. It would in this regard be of interest to further explore whether cells in S phase respond weaker to transcriptional stimulations as a general strategy to temporarily decouple DNA replication and transcription^54^.

Finally, we demonstrated that Live-seq and conventional scRNA-seq data can be seamlessly integrated with the aim of directly measuring a cell’s trajectory (**Figure 4l**). Indeed, while scRNA-seq provides the accuracy to resolve cellular heterogeneity, Live-seq experimentally maps cell state dynamics. As shown in this study, Live-seq can directly assign cell trajectories by assaying the same cells twice, here, before and after LPS stimulation. Integrating the transcriptomes of these reference cells with scRNA-seq data therefore provides a means to assign trajectories for cells that share the reference transcriptome signature. For such a hybrid approach, the monitoring of relatively few cells with Live-seq proved sufficient, which is readily possible with the current technology. Nonetheless, we expect the next generation of the Live-seq approach to allow for sampling of many more cells, including sequentially probed ones. This will in turn provide a future avenue to address some of the long-standing biological questions pertaining to cell dynamics or cellular phenotypic variation^8^.

In sum, Live-seq breaks new ground by enabling single cell transcriptome profiling as well as downstream analyses on the same cell at distinct time points (**Figure 4**). We anticipate that Live-seq will transform single cell transcriptomics from the current end-point type assay into a temporal analysis workflow.

## Methods

### Cell lines and culture

RAW264.7 and Hela cells were obtained from ATCC. RAW264.7 cells with *Tnf*-mCherry reporter and *relA*-GFP fusion protein (RAW-G9 clone) were kindly provided by Dr. Iain D.C. Fraser (NIH). The IBA cell line derived from the stromal vascular fraction (SVF) of interscapular brown adipose tissue of young male mice (C57BL6/J) was kindly provided by Prof. Christian Wolfrum’s laboratory (ETHZ). Primary ASPCs were isolated from subcutaneous fat tissue of C57BL/6J mice as previously described^55^. All these cells were cultured in high-glucose DMEM medium (Life Technologies, Carlsbad, CA) supplemented with 10% fetal bovine serum (FBS) (Gibco), and 1× penicillin/streptomycin solution (Life Technologies) in a 5% CO2 humidified atmosphere at 37 °C and maintained at less than 80% confluence before passaging.

#### Reagents

LPS (#L4391-1MG, Sigma, MO) was prepared in PBS at 100 μg/ml as a stock solution.

#### Oligos

Biotinylated-olig0-dT30VN (IDT):

/5Biosg/AAGCAGTGGTATCAACGCAGAGTACTTTTTTTTTTTTTTTTTTTTTTTTTTTTTTVN

TSO (Exiqon): AAGCAGTGGTATCAACGCAGAGTACATrGrG+G

Iso-TSO (Exiqon): iCiGiCAAGCAGTGGTATCAACGCAGAGTrGrG+G

Biotinylated-TSO (Exiqon): /5Biosg/AAGCAGTGGTATCAACGCAGAGTACATrGrG+G

Hairpin-TSO (Exiqon):

GGCGGCGCGCGCGCGCCGCCAAGCAGTGGTATCAACGCAGAGTACATrGrG+G

Biotinylated-ISPCR oligo (IDT): /5Biosg/AAGCAGTGGTATCAACGCAGAGT

Hairpin-TSO was annealed in IDT Duplex Buffer before use.

#### FluidFM setup

A FluidFM system composed of a FlexAFM-NIR scan head and a C3000 controller driven by the EasyScan2 software (Nanosurf), and a digital pressure controller unit (ranging from −800 to + 1000 mbar) operated by a digital controller software (Cytosurge) was used. A syringe pressure kit with a three-way valve (Cytosurge) was used in addition to the digital pressure controller to apply under- and overpressure differences larger than −800 and +1000 mbar. FluidFM Rapid Prototyping probes with a microchannel height of 800 nm were obtained from Cytosurge. A 400 nm wide triangular aperture was custom-milled by a focused ion beam near the apex of the pyramidal tips and imaged by scanning electron microscopy as previously described^56^. The FluidFM probes were plasma treated for 1 min (Plasma Cleaner PDG-32G, Harrick Plasma) and coated overnight with SL2 Sigmacote (Sigma-Aldrich) in a vacuum desiccator^56^.

### Optical microscopy setup

The FluidFM scan head was mounted on an inverted AxioObserver microscope equipped with a temperature-controlled incubation chamber (Zeiss), and coupled to a spinning disc confocal microscope (Visitron) with a Yokogawa CSU-W1 confocal unit and an EMCCD camera system (Andor). Phase-contrast and fluorescence images were acquired using 10× and 40× (0.6 na) objectives and a 2× lens switcher using VisiView software (Visitron). Microscopy images were analyzed using the AxioVision and ImageJ softwares.

### Enhanced Smart-seq2

The overall workflow follows the same steps as Smart-seq2^32^, but several modifications were introduced. RNA, Live-seq samples or single cells were transferred into a PCR tube with 4.2 μl lysis buffer, which contains 1μl dNTP(10mM each), 1 μl biotinylated-oligo-dT30VN oligo (10 μM), 2 μl 0.2% TritonX-100, 0.1 μl Recombinant ribonuclease inhibitor (40 U/μl, #2313, Takara), and 0.1 μl ERCC RNA Spike-In Mix (10^7^ dilution, #4456740, ThermoFisher). After briefly spinning, the PCR tubes/plates were heated at 72 °C for 3 min and cooled down on ice for 1 min. Then, 5.2 μl of reverse transcription mix (1.29 μl H_2_O, 0.06 μl MgCl_2_ (1M), 2 μl Betaine (5M), 0.08 μl biotinylated-TSO oligo (100 μM) (or other TSOs as listed in **Supplementary Figure 1d**), 2 μl Maxima H Mimus RT buffer (5X), 0.25 μl Recombinant ribonuclease inhibitor, 0.1 μl Maxima H Minus Reverse Transcriptase (200 U/μl, EP0751, ThermoFisher)) were added. The reverse transcription step was performed in a thermal cycler using the following program: 1) 42 °C for 90 min; 2) 50 °C for 2 min; 3) 42 °C for 2 min, go to step 2 for 4 cycles; 4) 85 °C for 5 min; 5) End. The reverse transcription products were directly used for PCR by adding 15 μl PCR mixture, which contains 12.5 μl KAPA HiFi HotStart ReadyMix (2X, 07958935001, KAPA), 0.25 μl Biotinylated-ISPCR oligo (10 μM) and 2.25 μl H_2_O. The PCR program involves the following steps: 1) 98 °C for 3 min; 2) 98 °C for 20 sec; 3) 67 °C for 15 sec; 4) 72 °C for 6 min, go to step 2 for X cycles (for Live-seq, X = 24; for scRNA-seq, X = 18); 5) 72 °C for 5 min; 6) End. The PCR products were purified twice with AMPure XP beads (A63882, Beckman Coulter) according to the manufacturer’s instructions with 0.6 volume of beads per 1 volume of PCR product. The concentration and profile of the purified DNA were measured by Qubit dsDNA HS Assay Kit (#Q32854, ThermoFisher) and fragment analyzer (Advanced Analytical), respectively. 1 ng of DNA was used for tagmentation in the following reaction: 1 μl TAPS buffer (50 mM TAPS-NaH, pH8.3, 25 mM MgCl_2_), 0.5 μl Tn5 (made in-house, 12.5 μM), H_2_O to 5 μl in total. The tagmentation reaction was assembled on ice, initiated by placing the mix on a thermal cycler at 55 °C for 8 min after which it was kept on ice. Then 1.5 μl SDS (0.2%) was added to inactivate the Tn5 enzyme. PCR reagent mixture was added directly to the tagmented DNA, which contains 1.5 μl dNTP (10 mM each), Nextera XT index primers (or other compatible indexing primers, with a final concentration of 300 nM for each primer), 10 μl KAPA HiFi Fidelity buffer (5X), 1 μl KAPA HiFi DNA Polymerase (1 U/µL, KK2101, KAPA) and H_2_O to 50 μl in total. Then, the DNA was amplified using the following program: 1) 72 °C for 3 min; 2) 98 °C for 30 sec; 3) 98 °C for 10 sec; 4) 63 °C for 30 sec; 5) 72 °C for 1 min, go to step 3 for 10 cycles; 6) 72 °C for 5 min; 7) End. The PCR products were purified twice with AMPure XP beads (A63882, Beckman Coulter) according to the manufacturer’s instructions with 0.6 volume of beads per 1 volume of PCR product. The concentration and profile of the purified DNA were measured using a Qubit dsDNA HS Assay Kit and fragment analyzer, respectively, after which the libraries could be sent for sequencing using Illumina sequencers. We thereby noted that the “cDNA” yield from the negative control (0 pg input RNA) (**Supplementary Figure 1g**) was mainly derived from oligo sequences, since sequencing of the respective library revealed mainly reads with a very high A/T content, which poorly aligned to the mouse/human genome (**Supplementary Figure 1j-k)**.

### Cytoplasmic biopsies

For extraction experiments, the cells were seeded and cultured for at least 18h onto 50-mm tissue-culture-treated low µ-dishes (Ibidi). Shortly before the experiment, the culture medium was replaced with 5 mL of CO_2_-independent growth medium supplemented with 10 % FBS, 1× penicillin/streptomycin solution, and 2mM Glutamine. Sampling buffer for preloading was prepared by supplementing a 0.2% solution of Triton-X 100 in nuclease-free water with 2 U μl^−1^ recombinant RNase inhibitors (Clonetech). Lysis buffer for extract transfer was prepared as that used in the enhanced Smart-seq2 protocol. While cells were maintained at 37°C for time-lapse microscopy before and after extractions, the extraction procedures were all performed at room temperature.

The probe reservoir was loaded with 10 µL of mineral oil (Sigma-Aldrich) and a pressure of Δ + 1000 mbar was applied to flow the oil into the microchannel. Once the probe microchannel was filled, the probe was shortly immersed in nuclease-free water, and then kept in air with the residual water carefully blotted off the probe holder with a kimwipe tissue. A 1.0-μl drop of sampling buffer was deposited onto an AG480F AmpliGrid (LTF Labortechnik). The cantilever was introduced into the drop using the micrometer screws to displace the AFM. Once the cantilever was located inside the drop, underpressure (−800 mbar) was applied for the suction of ∼0.5 pL of sampling buffer into the probe. The cantilever was then withdrawn from the drop using the micrometer screws.

Next, the pre-loaded probe was immersed in the cell sample experimental medium, the cell to be extracted was visualized by light microscopy, and the tip of the FluidFM probe was placed above the cytoplasm of the selected cell. The tip of the probe was then inserted into the cytoplasm through a forward force spectroscopy routine driven by the Z-piezo. The probe was then maintained inside the cell at constant force (500 nN). Underpressure larger than −800 mbar was applied to aspirate the cellular content into the probe, whereby the harvested cytoplasmic fluid immediately mixed with the preloaded sampling buffer. The pressure-assisted flow of the intracellular content into the FluidFM probe was interrupted by switching the pressure back to zero. We collected cytoplasmic extracts of 1.1 pL on average, ranging from 0.1 up to 4.4 pL. The probe was then retracted out of the cell, shortly immersed in nuclease-free water, and then kept in air with the residual water carefully blotted off the probe holder with a kimwipe tissue.

A 1.0-μl drop of lysis buffer was deposited onto an AG480F AmpliGrid (LTF Labortechnik). The cantilever was introduced into the drop, and overpressure (more than 1,000 mbar) was applied to release the extract. The microchannel was then rinsed three times by suction and release of lysis buffer into the probe. The 1.0-μl drop was then pipetted into a PCR tube containing an additional 3.2 μL of lysis buffer, and the solution was briefly centrifuged and stored at −80°C until further processing. The entire procedure took approximatively 15 min per extraction, and all steps were monitored in real time by optical microscopy in brightfield.

For the alternated human/mouse cell sampling, HeLa and IBA cells were seeded on separate dishes, after which cells were extracted alternatively from one or the other cell type as described above, using the same FluidFM probe.

LPS-stimulated RAW cells were extracted between 4 and 5 hours after the addition of LPS. For the sequential sampling of the same RAW cell, up to 14 cells were extracted as described above, with all the cells monitored within one vision field. After the first extractions, the cells were incubated at 37°C under time-lapse monitoring with intervals of 5 min. After 30 min, LPS was added to the dish for stimulation, and the cells were further monitored with 5 min intervals for another 30 min. The time-lapse intervals were then increased to 30 min, and the cells were further monitored for 3 hours. The temperature was then switched back to room temperature, and the same cells were extracted a second time, in a time interval of 4 to 5 hours after the addition of LPS.

### LPS stimulation of RAW cells

For stimulation of RAW cells, LPS solution was added to the 5 mL CO_2_-independent medium to reach a final concentration of 100 ng/mL.

### Determination of the pre-loaded and extracted volumes

For each extraction, the volume of preloaded sampling buffer and the volume of cytoplasmic extract mixed with the sampling buffer were measured on brightfield micrographs using the AxioVision software. The area occupied by the aqueous solutions confined in the cantilever was multiplied by the channel height of 0.8 μm, and the volume of the hollow pyramidal tip (90 fL) was added. To determine the volume of extracted cytoplasmic fluid, the volume of preloaded sampling buffer was subtracted to the volume of mixed sampling buffer and extract.

### Determination of the cell volumes

To quantify the whole cell volumes, IBA and ASPC cells were dissociated by trypsinization and RAW264.7 cells with a cell scraper. The dissociated cells were then imaged in brightfield with the 40× objective and the 2× lens switcher, and the diameter of the rounded cells was measured from the micrographs using the AxioVision software. Three diameters were measured and averaged for each cell (N=277 for ASPC and N=500 for IBA and RAW). The volumes were calculated using the formula for a sphere. For the longitudinal measurements of RAW cell volumes, the areas of the semi-adherent cells were measured on brightfield images acquired with the 40× objective and the 2× lens switcher at multiple time points before and after extraction, and cell volumes were calculated assuming a spherical cell shape.

### Time-lapse monitoring of mCherry expression

For live imaging of RAW-G9 cells expressing mCherry, the ibidi dish was covered and the temperature was maintained at 37°C. Time-lapse images were acquired at 5 min intervals for 1 h, then at 30 min intervals overnight. For each time point, two sequential frames were acquired, in brightfield and for mCherry (561 nm laser and 609/54 nm emission filter), using the 40× objective and the 2× lens switcher. At the end of the time-lapse recording, 2 µL of tricolor calibration beads (Invitrogen™ MultiSpeck™ Multispectral Fluorescence Microscopy Standards Kit, Life Technologies Europe B.V.) was added to the sample, and at least 30 individual beads were imaged with the 561 laser. For image quantification, all the cells that moved out of view or focus, died, or overlapped with other cells were excluded from analysis in each time-lapse frame. Cell boundaries were all manually defined in Fiji, and background intensities were measured for each time point and subtracted from all intensity values. We analysed 77 extracted cells, 122 LPS-stimulated cells that were not extracted, and 23 cells that were not extracted and not stimulated with LPS. The fluorescence intensity of the calibration beads with subtracted background intensity was measured using Fiji, and the average of the 30 bead intensities was used to normalize the fluorescence measurements of the cells acquired in different experiments. The area under a curve (AUC) was calculated from 3 hour to 7.5 hours post LPS treatment. A two-sided Wilcoxon rank-sum test was used to examine the differences between conditions.

### Cell viability after extraction

To evaluate the post-extraction viability of ASPC (N=33) and IBA (N=37) cells, the cells were stained between 2 and 4 hours after extraction using a LIVE/DEAD^®^ Cell Imaging Kit 488/570 (Invitrogen), and following the manufacturer’s protocol. To evaluate the post-extraction viability of RAW264.7 cells, the extracted cells (N= 72) were monitored by time-lapse microscopy during ∼10 hours at 30 min intervals. Cells were evaluated as dead or alive based on their morphology, movements, and expression of mCherry in response to LPS stimulation. The post-extraction viability was assessed for 42 cells extracted once before stimulation, 30 cells extracted once after LPS stimulation, and 10 cells extracted twice, before and after stimulation. The viability of all the cell types was calculated as an absolute value without normalization.

### scRNA-seq

Cells were trypsinized and dissociated into a single cell suspension. After passing through a 40 μm cell constrainer and DAPI staining, live singlet cells were sorted into 96 well PCR plates with 4.2 μl lysis buffer (mentioned above). At least 3 wells without cells were preserved as negative controls. The plate was quickly spun and further processed using the Smart-seq2 method^32^.

To perform scRNA-seq on cells post Live-seq extraction, cells were first cultured in a dish containing a silicone micro-Insert (ibidi, #80409) at a density of around 20 cells/well. The insert was then removed just before Live-seq sampling and all the cells were subjected to Live-seq extraction as mentioned above. In order to extract all the cells within a reasonable time window (1 hour in this experiment), the extracted cytoplasm was not preserved. 1 hour and 4 hours post extraction, the cells, along with not extracted control cells, were harvested and single cells were picked using a serial dilution approach. The downstream processing followed a similar workflow as the Smart-seq2 method.

### Cell cycle analyses

The cell cycle reporter vector pCSII-EF-miRFP709-hCdt(1/100) was a kind gift from Vladislav Verkhusha (Addgene plasmid # 80007; http://n2t.net/addgene:80007; RRID:Addgene_80007). Lentivirus carrying this vector was transduced into RAW-G9 cells after which miRFP709-positive cells were sorted to enrich for transduced cells. The cells were seeded on an Ibidi dish and monitored for miRFP709 (640 nm laser and 700/75 nm emission filter) and in brightfield for 24 h at 60 min intervals with the 40× objective and the 2× lens switcher. The experimental growth medium was then exchanged, LPS was added at a final concentration of 100 ng/mL, and the cells were monitored for another 24 h at 60 min intervals in brightfield, for miRFP709, and for mCherry (561 nm laser and 609/54 nm emission filter). Fluorescence intensities of the cells were measured as described above (**Time-lapse monitoring of mCherry expression)** for both fluorescent reporters. The mCherry intensities were further normalized using the beads calibration.

### Live-seq and scRNA-seq data analysis

#### Dataset description

There are 10 IBA cells and Hela cells processed through Live-seq sampling in alternative fashion. In addition, 443 Live-seq samples were prepared across four experimental replicates, with each of them containing both single and sequential sampling events. All the data from these 443 cells were used for the analyses presented in **Figures 2-4**. 343 cells were processed using conventional scRNA-seq as part of two experimental replicates and the data linked to these cells are presented in **Figures 2-4**. Finally, to evaluate the potential molecular perturbation of cytoplasmic sampling, scRNA-seq data of IBA cells 1 hour (49 cells) and 4 hours (43) post extraction was generated using cells (70 cell) that were not subjected to such sampling as control.

#### Alignment and feature counting

Libraries were sequenced in either 75 single-read or 2x 75 paired-end format using Nextera indexes on Illumina Nextseq500 or Hiseq4000 sequencers at the Gene Expression core facility (EPFL). To keep consistency, only read1 with 75 bp was used for further analysis. The reads were aligned to the human (hg19/ GRCh38) or mouse (mm10/GRCm38) genomes using STAR (2.6.1c)^57^ with default settings and filtered for uniquely mapped reads. Then, the number of reads per feature (gene) was counted using HTseq (0.10.0)^58^ with parameter “htseq-count -s no -m union -f bam” and the gene annotation of Ensembl release 87 supplemented with ERCC, EGFP and mCherry features was used. The counts of all samples were merged into a single gene expression matrix, with genes in rows and samples in columns (Gene Expression Omnibus (GEO) accession number GSE141064, processed data).

To evaluate the cross-sample contamination, we sampled human and mouse cells alternatively. Reads were aligned to the mixed human: mouse reference genome (hg38 and mm10) using STAR and the number of reads per feature was counted by HTseq using the same settings as mentioned above. Digital gene expression matrices were generated for each species. Downstream data analysis was performed using R (version 3.5.0), plots generated using the R package ggplot2 (version 3.2.1).

#### Cell and gene filtering

ERCCs were used for technical evaluation by testing the correlation with the expected number of spike-in RNA molecules, and not included for further analysis. The top 20 genes found in negative controls (samples were merged) are likely due to misalignment/sequencing errors of the oligos (**Supplementary Table 5**), and were thus also removed from downstream analyses. Ribosomal protein coding genes were removed as they confound the downstream differential gene expression analysis, consistent with previous findings^59^. The downstream analysis followed the procedures of the Seurat R package (v3.0). Samples showing low quality were filtered out, with quality cutoffs being: i) the number of genes <1000, ii) the mitochondrial read ratio >30%, or iii) the uniquely mapped rate <30%. Then, the data was normalized to the total expression, multiplied by a scale factor of 10,000, after which a pseudo count was added and the data log transformed.

#### Feature selection, dimensionality, reduction clustering and others

The top 500 highly variable genes were chosen based on the variance stabilizing transformation (vst) result (function: FindVariableFeatures(object, selection.method = “vst”)). Scaling was applied to all the genes using the function “ScaleData”. The scaled data of the 500 highly variable genes was used for PCA analysis. The first 10 principal components (PCs) were chosen for further clustering and tSNE analysis, based on the ranking of the percentage of variance explained by each PC (ElbowPlot function). Both the Seurat-embedded graph-based clustering and independent hierarchical clustering (ward.D2 method) approaches were applied, and both methods yielded similar results in our analysis. We noted though that RAW_LPS cells split into two clusters due to a batch effect (**Figure 2b** and **Supplementary Figure 2f**), which is why they were merged together manually. For data visualization, individual cells were projected on the basis of the 10 PC scores onto a two-dimensional map using t-SNE^60^.

To evaluate the effect of varying sequencing depth on the clustering of Live-seq data, we down-sampled the raw counts to the desired number. The down-sampled matrices were analyzed in the same way as described above. The clusters were consistent with the original analysis, indicating that the clusters were not driven by sequencing depth. A two-sided Wilcoxon rank-sum test method was used to find the differentially expressed genes (both positive and negative) among clusters, with a fold-change cutoff at 0.25 (log-scale). The pseudo-genes were filtered out^61^. The top 100 marker genes were loaded into the EnrichR package^37^ to determine gene enrichment among the biological processes of Gene Ontology and predict the cell types using the Mouse Gene Atlas database. GSEA was performed in R using the ClusterProfiler package^62^ version 3.14.3 and msigdbr version 7.1.1 using default settings with all the gene expression changes. Both wikipathways-20200810-gmt-Mus_musculus.gmt and the GO terms obtained from the R database org.Mm.eg.db (version 3.10.0) were used as reference.

The scores of cell cycle phases were calculated using the Seurat function “CellCycleScoring”, based on canonical markers^63^.

To integrate Live-seq and scRNA-seq data, the canonical correlation analysis (CCA) and mutual nearest neighbors (MNNs)-based approaches embedded in Seurat v3 were used. Specifically, the top 500 highly variable genes of both datasets were chosen for PCA analysis independently. The first 10 PCs of each were used to identify the anchors and for data integration. As another level of control, data stemming from cells belonging to each cluster in both Live-seq and scRNA-seq analyses were collapsed into a “bulk” RNA-seq dataset after which the Pearson’s correlation between the “bulk” Live-seq and the “bulk” scRNA-seq datasets was determined. To evaluate the effect of varying sequencing depth on data integration, we down-sampled the raw counts to the desired number. The down-sampled matrices were then analyzed in the same way as described above.

#### Live-seq and live cell imaging integration

Among the 40 cells that were both subjected to Live-seq and tracked for LPS-induced *Tnf*-mCherry fluorescence, 17 of them passed the quality control as mentioned in the Live-seq section. For each cell within the time-course, we calculated the intercept (i.e. basal expression) and slope (i.e. extent of response) using a linear model between the time after LPS treatment and the natural log *Tnf*-mCherry fluorescence. As we were mainly interested in the initial response and since the curve is linear from the first 3 to 7.5 hours (**Figure 4c** and **Supplementary Figure 4a**), we only used values from this time window. To rank genes, we then constructed a linear model that predicts the intercept or slope based on the expression of a gene measured by Live-seq prior to treatment with LPS. Given the limited number of cells and thus to increase the statistical power of the models, only the 500 most variable genes (instead of all genes) from both the RAW-G9 Live-seq and scRNA-seq data were used, with genes not detected in Live-seq data removed. The genes were then ranked based on their respective R^2^ values.

### Trajectory interference

To infer the trajectory from conventional scRNA-seq data, the dynverse R package^24^ with multiple wrapped trajectory inference methods was used^17, 19, 64–66^. The parameters shown below were used for the “answer_questions” function-“multiple_disconnected = NULL, expect_topology = NULL, expected_topology = NULL, n_cells = 333, n_features = 1000, memory = “100GB”, docker = TRUE”. The most suggested methods were chosen based on the guidelines provided by dynverse. In addition, we also applied the Monocle DDRTree method ^18^ given its widespread use. All these methods were run in the docker with default parameters. For the RNA velocity analysis, annotated spliced, unspliced and spanning reads in the measured cells were generated in a single loom file using the command line “velocyto run_smartseq2 -d 1” function. This also generates an HDF5 file containing detailed molecular mapping information which was used for the analysis model based on gene structure. Three different algorithms were used following the velocyte pipeline (http://pklab.med.harvard.edu/velocyto/notebooks/R/chromaffin2.nb.html): 1) cell kNN pooling with the gamma fit based on extreme quantiles, 2) relative gamma fit, without cell kNN smoothing, and 3) velocity estimate based on gene structure.

### Analysis of the most variable genes from primary macrophage scRNA-seq data

We retrieved single-cell expression data of 5 macrophage subsets from the GEO database (GSE117081)^67^. We calculated for each gene a standardized variance by modelling the relationship between the observed mean expression and variance using local polynomial regression, as implemented in the Seurat (3.1.4) FindVariableFeatures function. Genes of the KEGG NF-κB signaling pathway downstream TLR4 receptor were highlighted.

### Bioethics

All mouse experiments were conducted in strict accordance with the Swiss law, and all experiments were approved by the ethics commission of the state veterinary office (60/2012, 43/2011).

### Data availability

All Live-seq and scRNA-seq data are available in the Gene Expression Omnibus (GEO) with accession number GSE141064.

### Code availability

The codes used for the analysis are incorporated into the Methods sections listed above.

## Acknowledgements

We thank Tomaso Zambelli, Julie Russeil and Petra C. Schwalie for support and Vincent Gardeux, Joern Pezoldt, Gioele La Manno, Judith Kribelbauer and Wouter Saelens for reviewing the manuscript and data analysis. We thank Dr. Iain D.C. Fraser (NIH) for kindly providing the RAW-G9 cells. We thank the Gene Expression (GECF, EPFL) and Flow Cytometry (FCCF, EPFL) core facilities for technical support. This work was supported by a Swiss National Science Foundation Grant (310030_182655), a Precision Health & related Technologies Grant (PHRT-502) and institutional funding (EPFL) to B.D., a Marie Skłodowska-Curie fellowship, EPFL Fellows (665667) to W.C., and by a grant from the Volkswagen foundation (Initiative “Life”), a European Research Council Advanced Grant (no. 883077) and institutional funding (ETH Zurich) to J.A.V.

## Author contribution

W.C., O.G.G, J.A.V and B.D. designed the study and wrote the manuscript. W.C., O.G.G performed experiments and data analysis with the support of R.D., P.Y.R., M.Z., C.G.G. All authors read and approved the final manuscript.

---

**Supplementary Figure 1. Relative to Figure 1.**
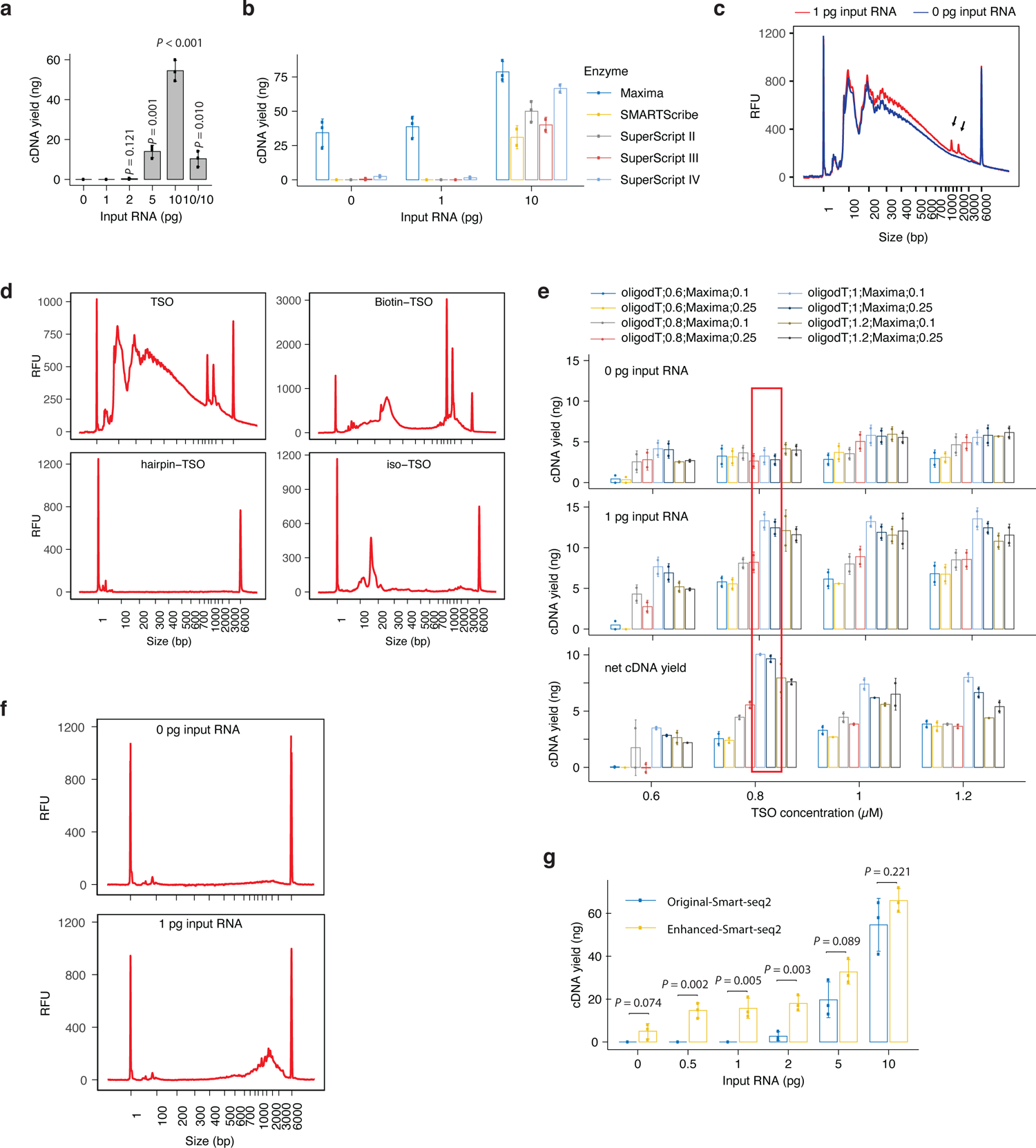
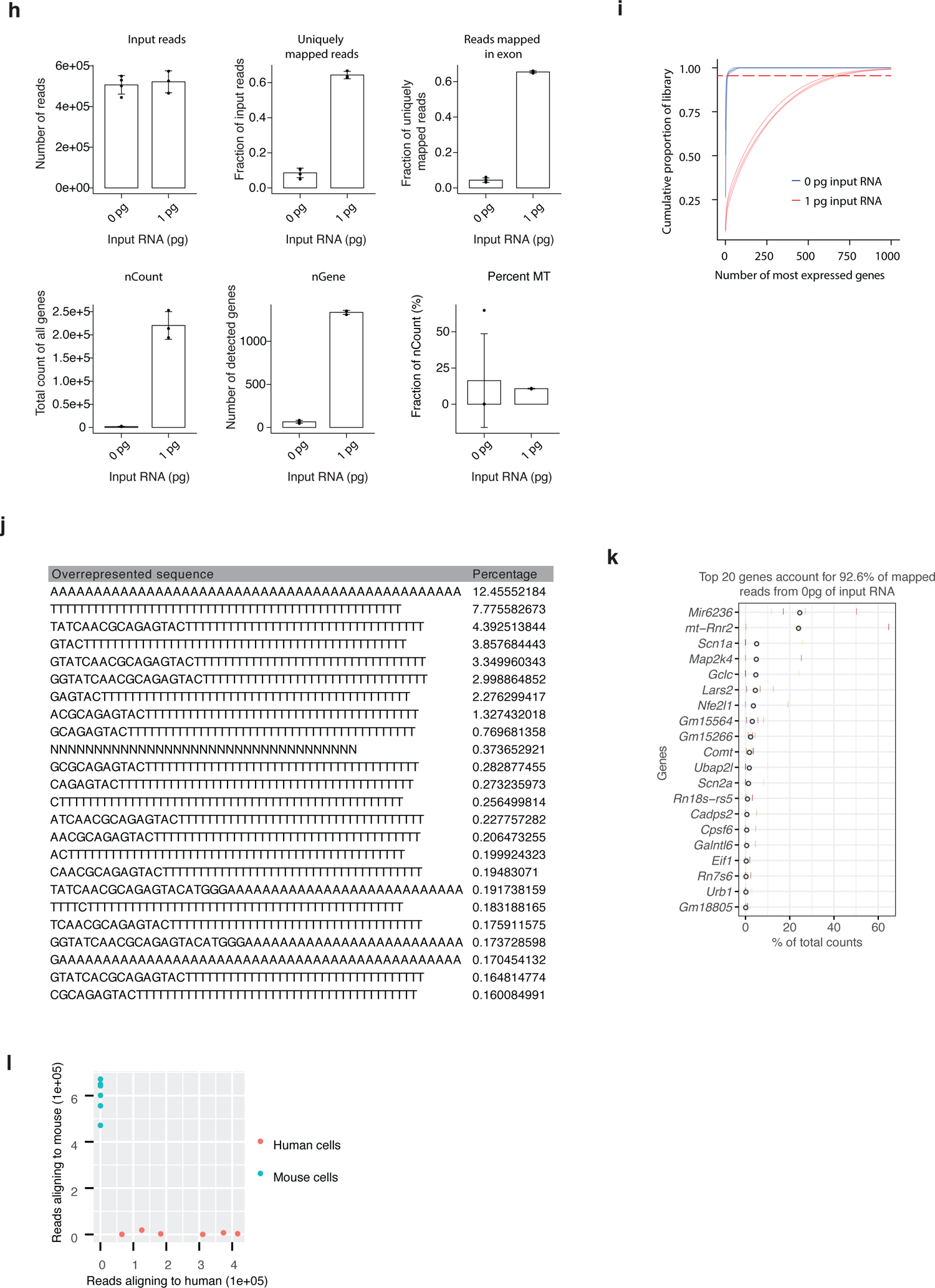
(**a**) cDNA yields of different amounts of input total RNA using the Smart-seq2 method. 10/10 represents the cDNA yield from 10% of the reverse transcribed material produced from 10 pg of total RNA that was used for PCR amplification. N = 3 replicates. *P* values were determined by two-sided *t*-tests, with each condition compared to 0 pg input RNA. (**b**) Screening of distinct reverse transcriptases, as listed. 0 (negative control), 1, and 10 pg of total RNA was used as input. N = 3 replicates. (**c**) The profile of cDNA using Maxima H-reverse transcriptase. The hedgehog pattern from 100 – 3000 bp (with peaks from 100 to 500 bp) reflects the presence of oligo concatemers. The arrows show two unique peaks in cDNA from 1 pg RNA compared to the negative control (0 pg input RNA), suggesting that bone fide signal is masked by concatemers. (**d**) Profiles of cDNAs that were generated using different 5’ modified TSO primers. Maxima H-reverse transcriptase was used in all conditions. The condition using Biotin-TSO shows a high cDNA yield between 500 to 3000 bp with a relatively low amount of oligo dimers or concatemers (the hedgehog-like pattern with peaks from 100 to 500 bp). (**e**) cDNA yields of distinct experimental set-ups involving different combinations of Biotin-TSO, oligo-dT, and reverse transcriptase from 0 pg input RNA (upper panel) and 1 pg input RNA (middle panel). The net cDNA yield was calculated as the yield from 1 pg of input RNA minus signal stemming from 0 pg of input RNA (lower panel). The workflow with the highest net yield is highlighted using a red rectangle and forthwith termed “enhanced Smart-seq2”. N = 3 replicates. (**f**) Profiles of cDNA derived from the enhanced Smart-seq2 protocol. (**g**) Enhanced Smart-seq2 is more sensitive compared to the original Smart-seq2 approach in the low input (0.5-2 pg) RNA range. The cDNA derived from 0 pg input RNA is derived from a few genes (see j, k). N = 3 replicates for all conditions. *P* value determined by a two-sided *t*-test. (**h**) Quality control of the enhanced Smart-seq2 workflow based on the parameters that are listed above each panel, comparing negative control (0 pg, N = 4 replicates) to IBA cell RNA (1 pg, N = 3 replicates). nGene: number of detected genes. nCount: total count of all genes. Percent MT: percentage of counts from mitochondrial genes. (**i**) The cumulative proportion of each library (y axis) assigned to the top-expressed genes (x axis). The top 20 genes absorb around 95% of all the reads in the negative control (0 pg input RNA, N = 4 replicates), while the ∼700 top genes take that same portion of reads in samples with 1 pg input RNA (N = 3 replicates). The dashed line indicates the 95% proportion. (**j**) Overview of the overrepresented sequences from the negative control. They are mostly sequences that are derived from the oligo-dT and TSO. (**k**) The proportion of reads mapped to each gene in negative control samples. The top 20 genes account for more than 90% of all reads. (**l**) Human (Hela) and mouse (IBA) cells were alternatively sampled by Live-seq using the same tips, with a washing step being incorporated between each sampling. The number of reads that mapped to respectively the human or mouse genome for each sample was determined to assess potential cross-sample contamination. The IBA samples are the same ones used for the analysis shown in Figure 1b. Error bars represent the mean +/- SD.

**Supplementary Figure 2. Relative to Figure 2.**
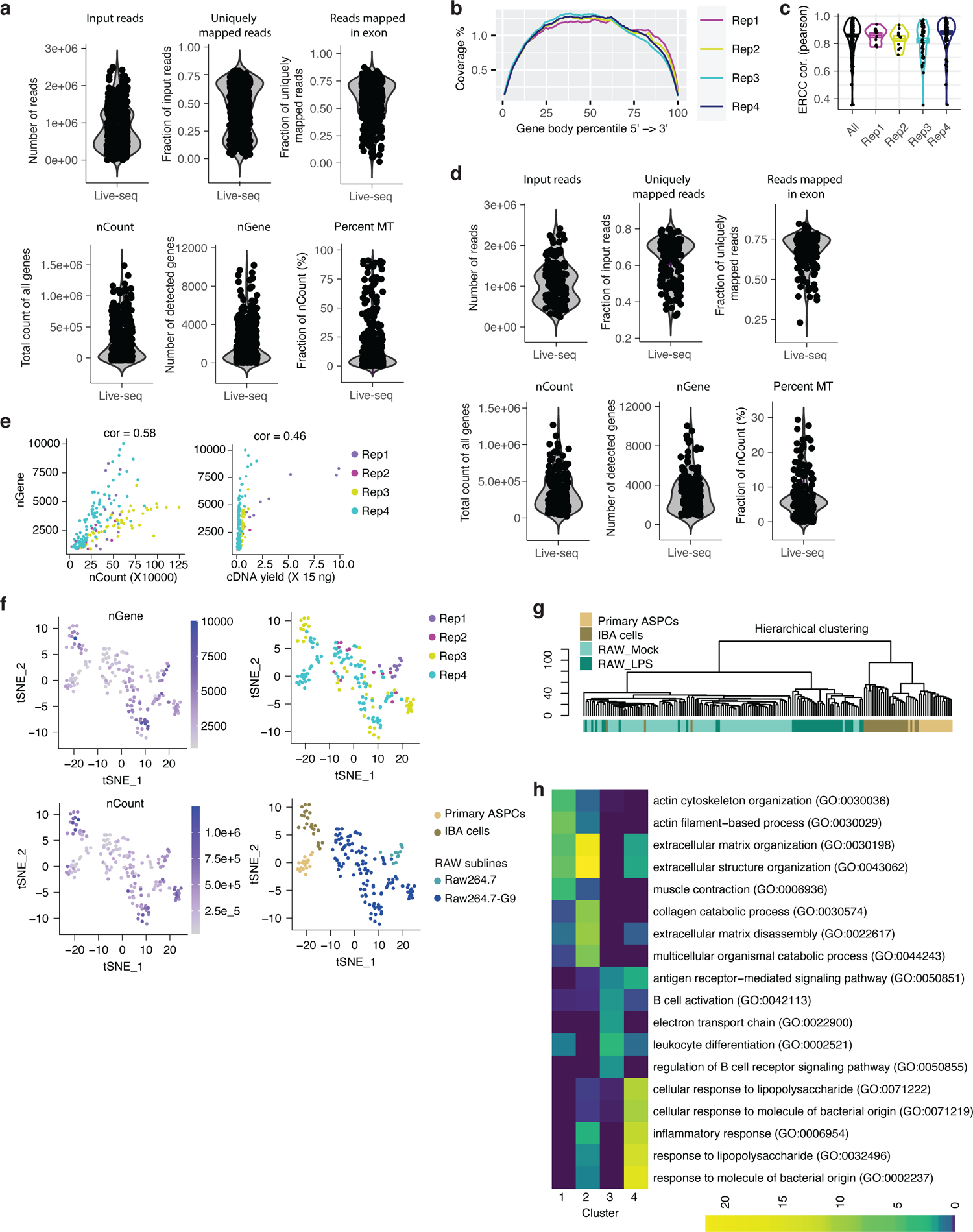
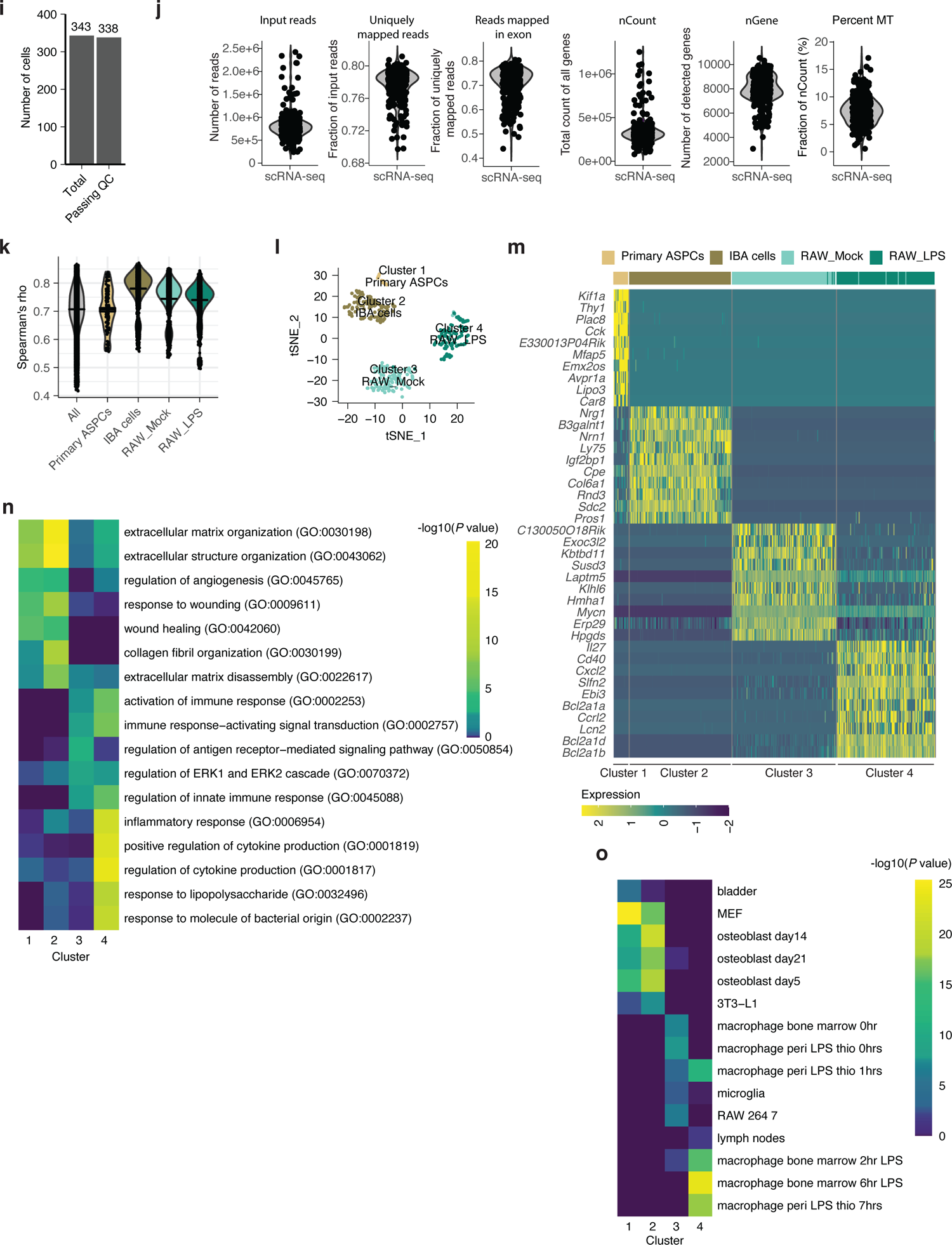
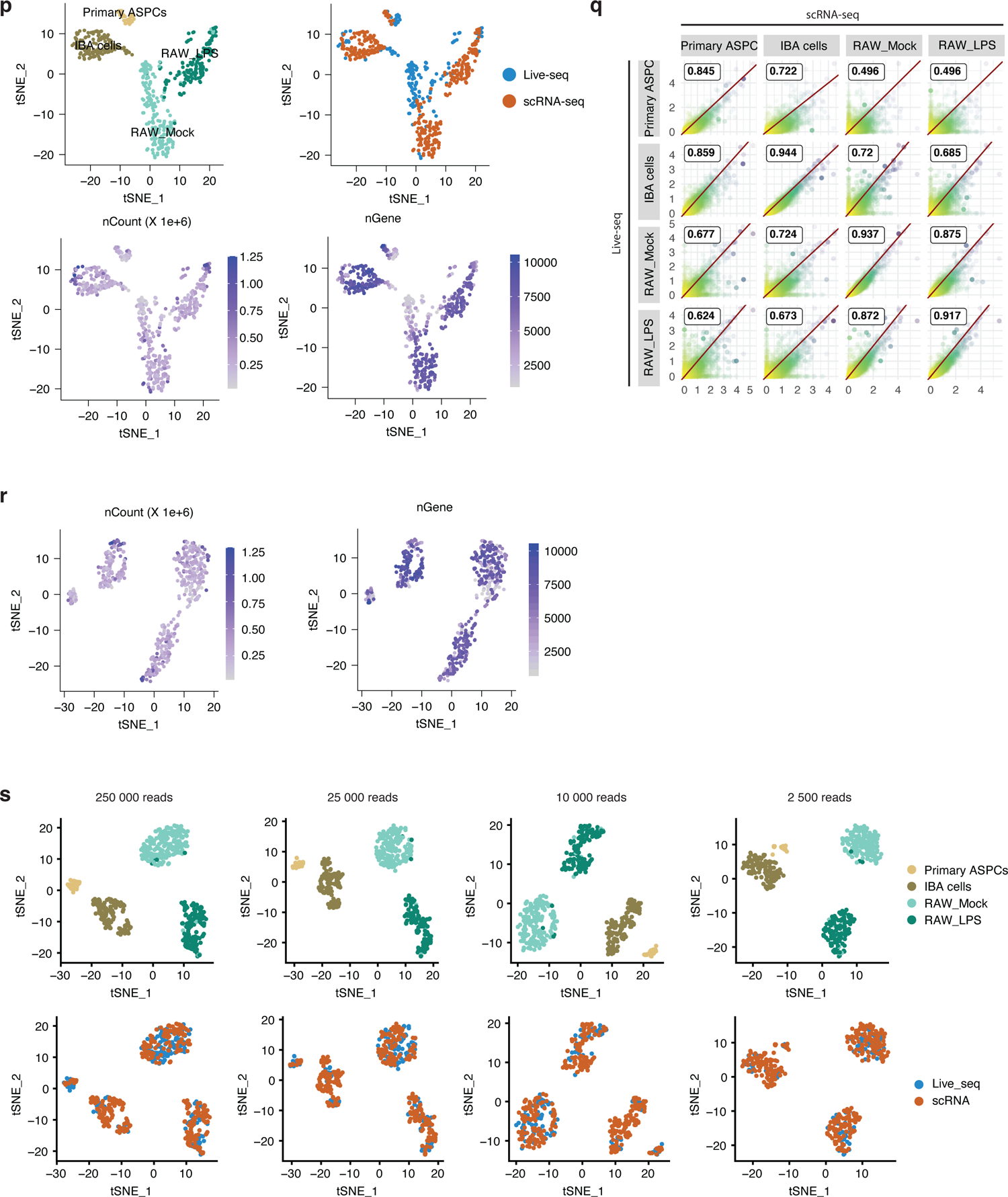
(**a**) Number of input reads (input reads), the rate of reads uniquely mapped to the genome (uniquely mapped reads), the fraction of reads mapped to exons (Reads mapped in exon), total counts of all genes (nCount), number of detected genes (nGene) and the percentage of counts from mitochondrial genes (percent MT), for all 390 Live-seq samples / libraries are shown. N = 4 replicates, a total of 390 cells. (**b**) The read distribution along the body of housekeeping genes for the 175 samples passing our quality control across different sample replicates (Rep1, Rep2, Rep3, Rep4). (**c**) Plotted Pearson correlation values between the amount of ERCC detected and expected for all 175 samples passing our quality control across different sample replicates. (**d**) Quality control of Live-seq samples similar to (a), but only showing those that passed the cutoff. N = 4 replicates, a total of 175 cells. (**e**) Plotted correlations between the number of detected genes on the one hand and respectively the total count of all genes (nCount, left panel) and the cDNA yield (right panel). The Pearson correlation value is displayed above each plot. N = 4 replicates, a total of 175 cells. (**f**) tSNE-based visualization of clusters, nGene, nCount, replicates, and cell lines. (**g**) Hierarchical clustering (ward.D2 method) of the 175 Live-seq samples based on the top 500 highly variable genes. (**h**) GO term enrichment of each cluster using the top 100 marker genes. (**i**) The number of cells processed with conventional scRNA-seq and those passing the quality control using similar parameters as those used for Live-seq cells. (**j**) Quality control of scRNA-seq data plotting distinct parameters as indicated above each panel. Similar to (a). (**k**) Gene expression correlation (Spearman’s rho) among all scRNA-seq-based transcriptomes stratified according to cell type and treatment. (**l**) tSNE-based scRNA-seq data visualization of the clusters using the top 500 highly variable genes. (**m**) Heatmap showing the top 10 differentially expressed genes stratified according to the four scRNA-seq clusters. (**n**) GO term enrichment analysis of the four scRNA-seq clusters using the top 100 differentially expressed genes. (**o**) Mouse gene atlas-based prediction of cell type/state of each cluster using the top 100 marker genes. (**p**) tSNE-based visualization of integrated scRNA-seq and Live-seq data according to cell type or treatment, approach, number of detected genes, and total counts of all genes without batch correction. (**q**) The correlation between simulated bulk data (i.e. based on the aggregation of each scRNA-seq or Live-seq expression profile into bulk-like data) across cell type and treatment. The Pearson correlation value is shown inside each of the subpanels. (**r**) Additional quality controls relative to Figure 2e: visualization of Live-seq and scRNA-seq data after anchor-based data integration (**Methods**) according to the number of detected genes and total gene expression count. (**s**) Evaluation of the data integration as in Figure 2e whereby each cell was downsampled to the indicated number of reads.

**Supplementary Figure 3. Relative to Figure 3.**
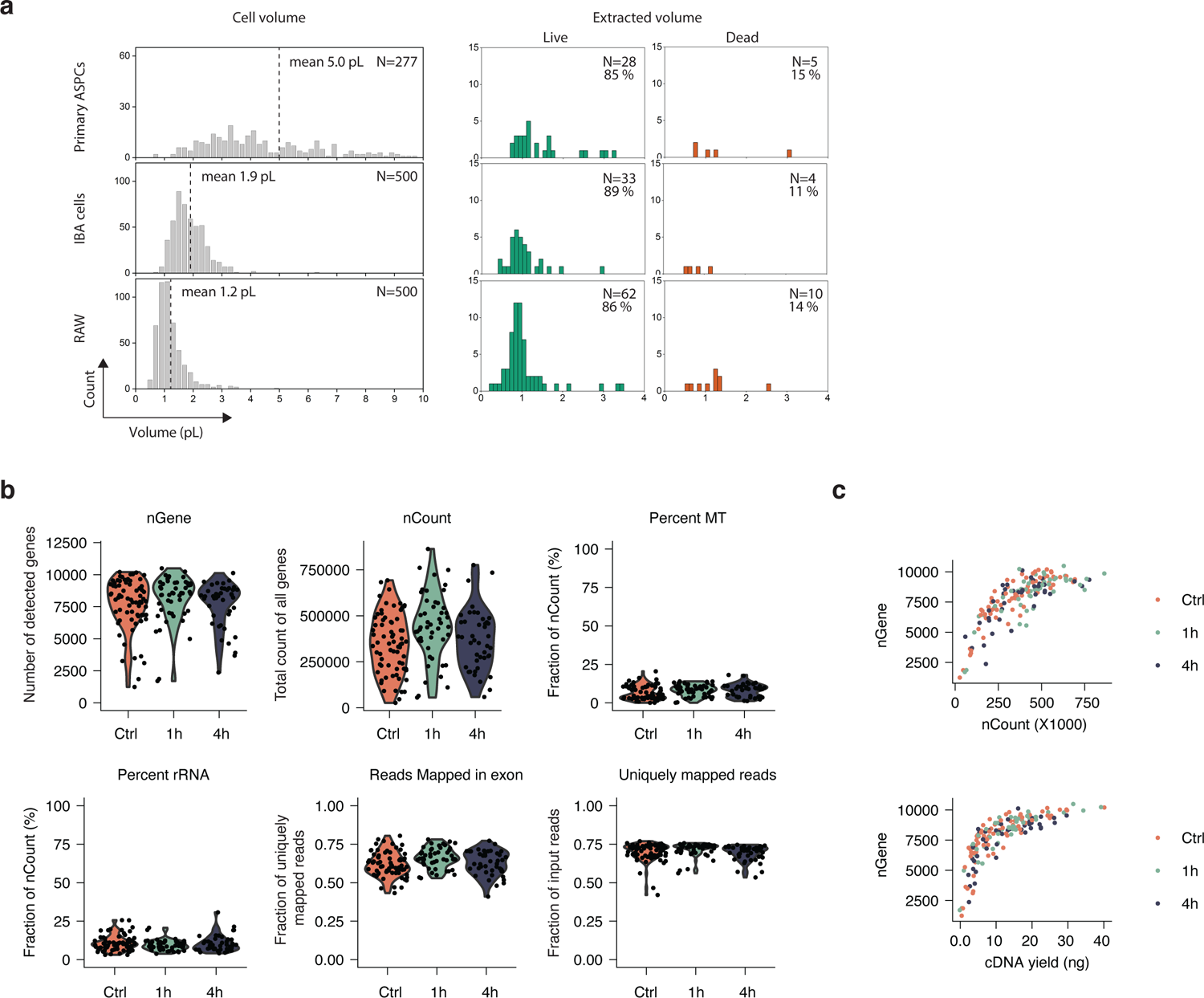
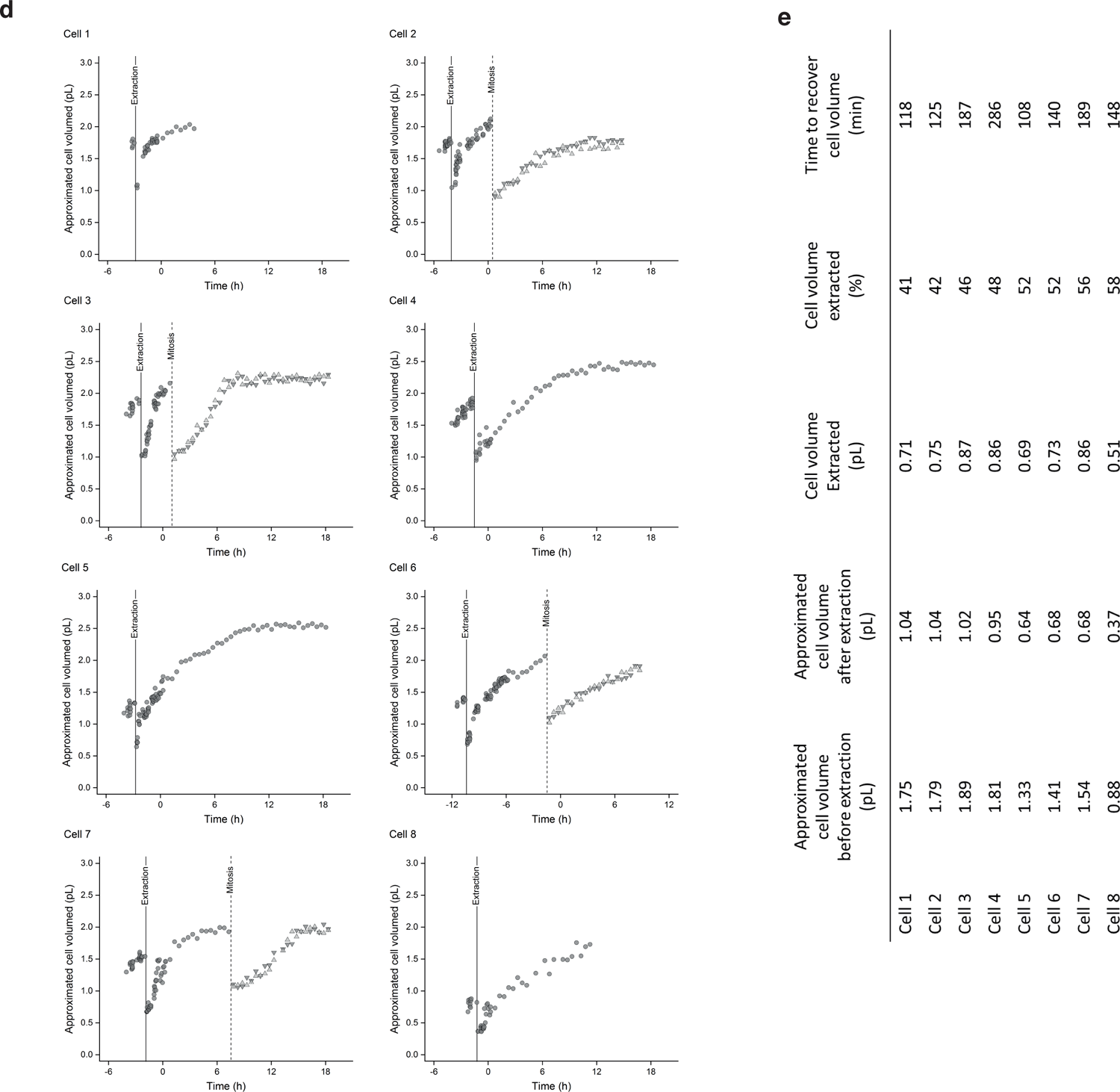
(**a**) Distribution of estimated cell volumes (left) and post-extraction viability as a function of the Live-seq sampled volumes (right) for primary ASPCs (N = 33), IBA (N= 37) and RAW cells (N= 72). The viability is an absolute value without normalization. (**b**) Quality control of the scRNA-seq data of the control cells, as well as cells 1 hour and 4 hours post Live-seq extraction, respectively, shown in Figure 3a. (**c**) Plotted correlations between the number of detected genes on the one hand and respectively the total count of all genes (nCount, upper panel) and the cDNA yield (lower panel), were undistinguishable between the respective cell categories. (**d**) Longitudinal measurements of RAW cell volumes. Cells were exposed to LPS at time 0. Vertical lines indicate the time of extraction (solid) and of mitosis (dashed). After mitosis, both daughter cells were monitored (upside and downside triangles). (**e**) The cell volume changes, extracted cytoplasm volumes, and recovery times were calculated based on the profiles in (d).

**Supplementary Figure 4. Relative to Figure 4.**
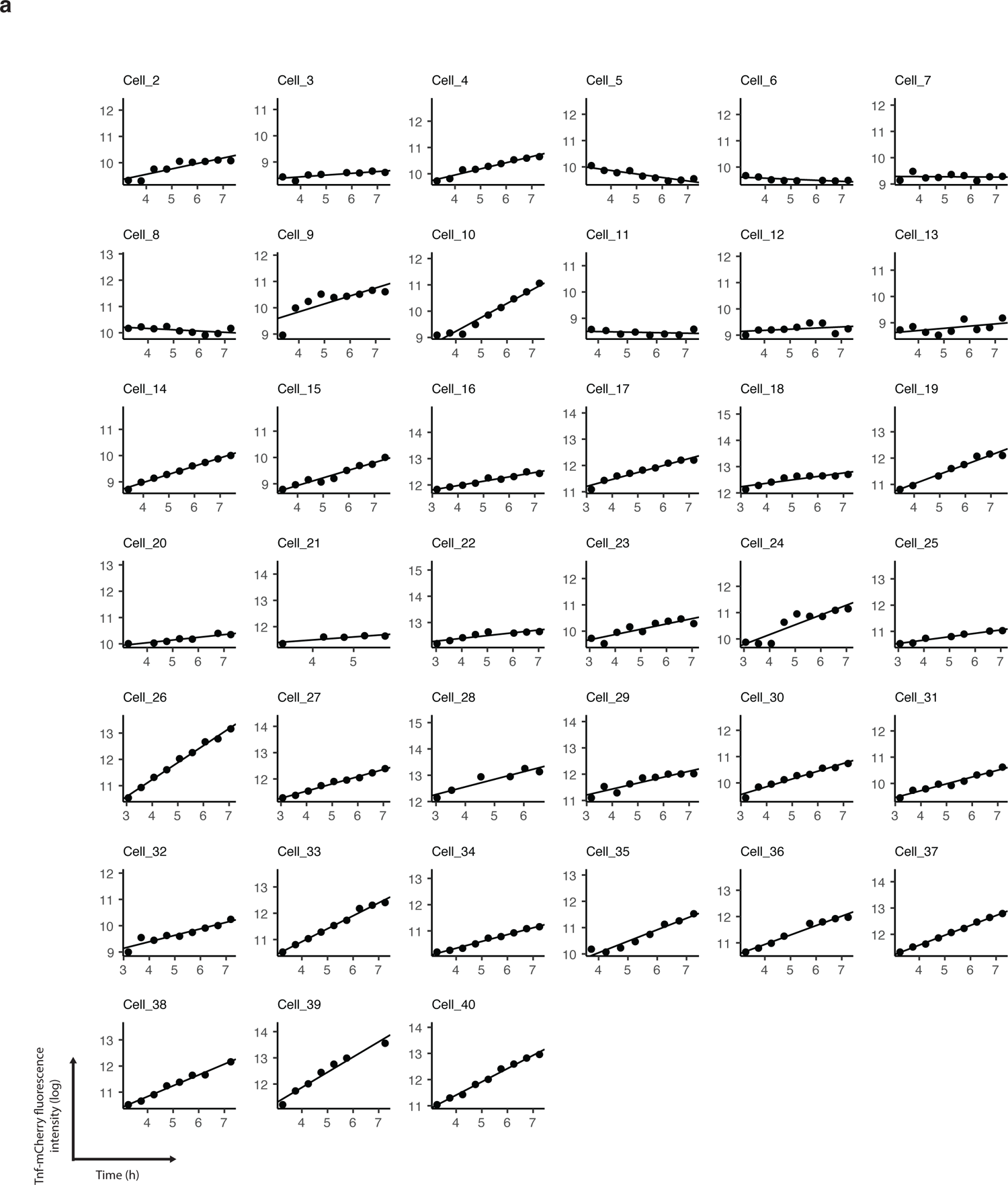
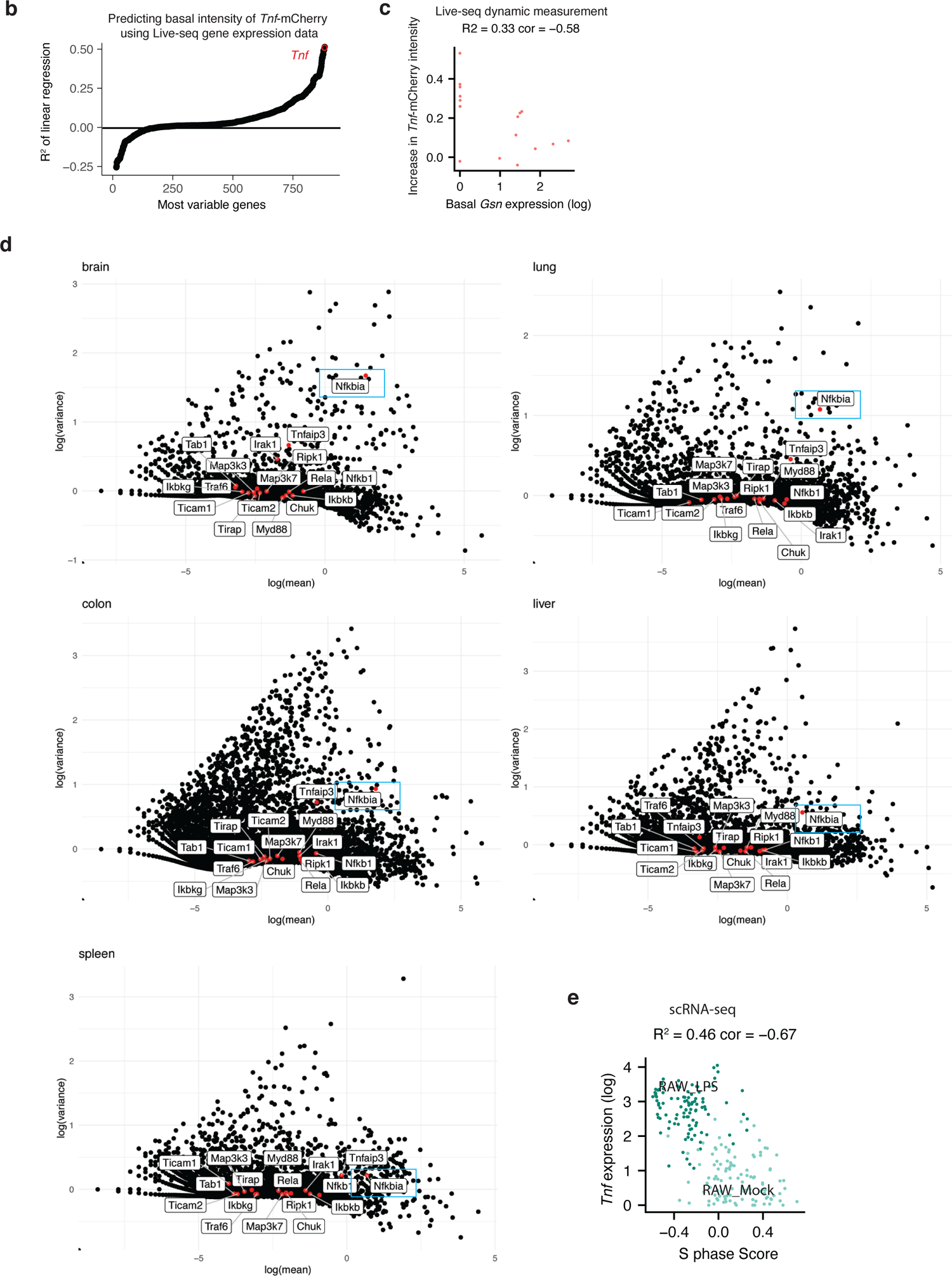
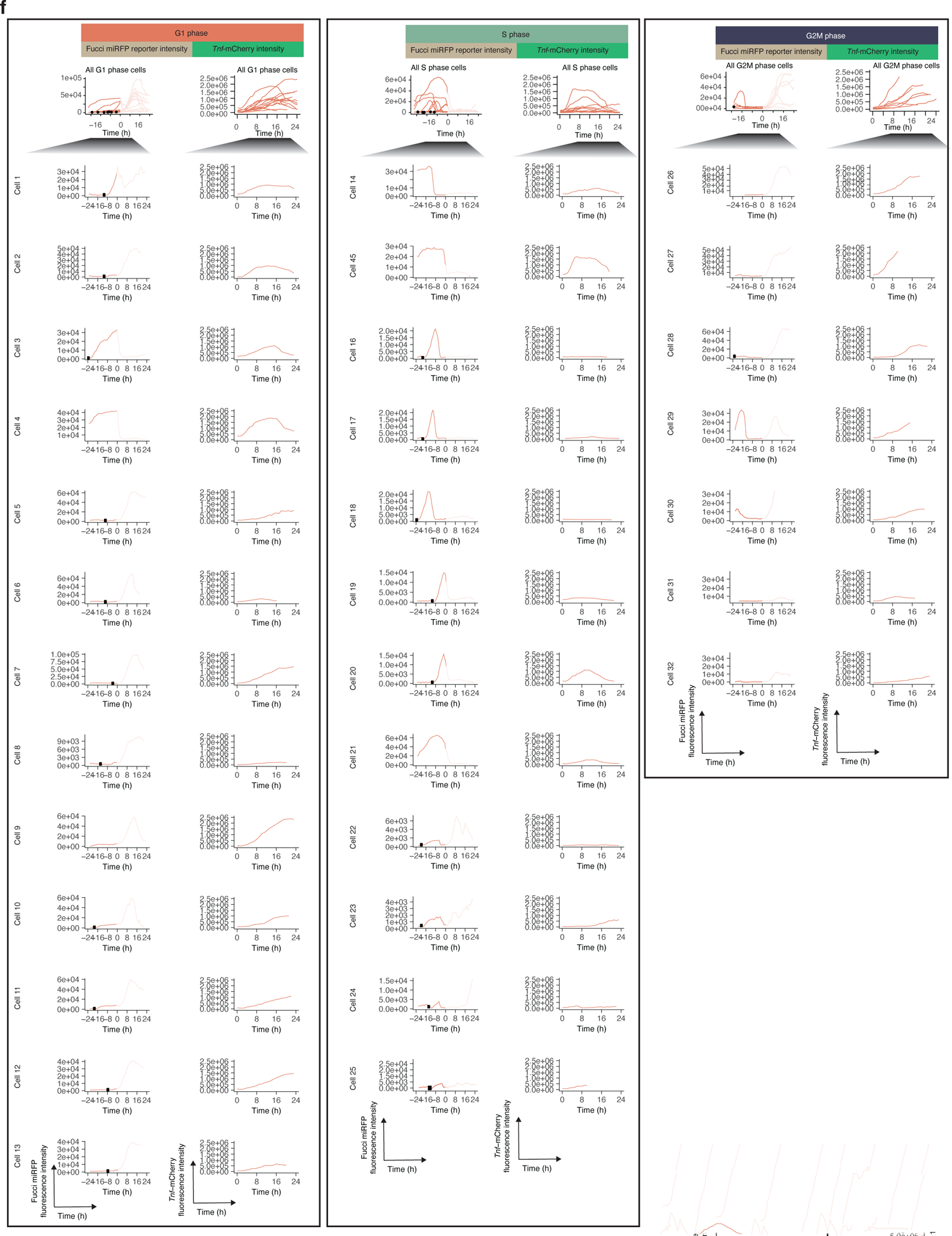
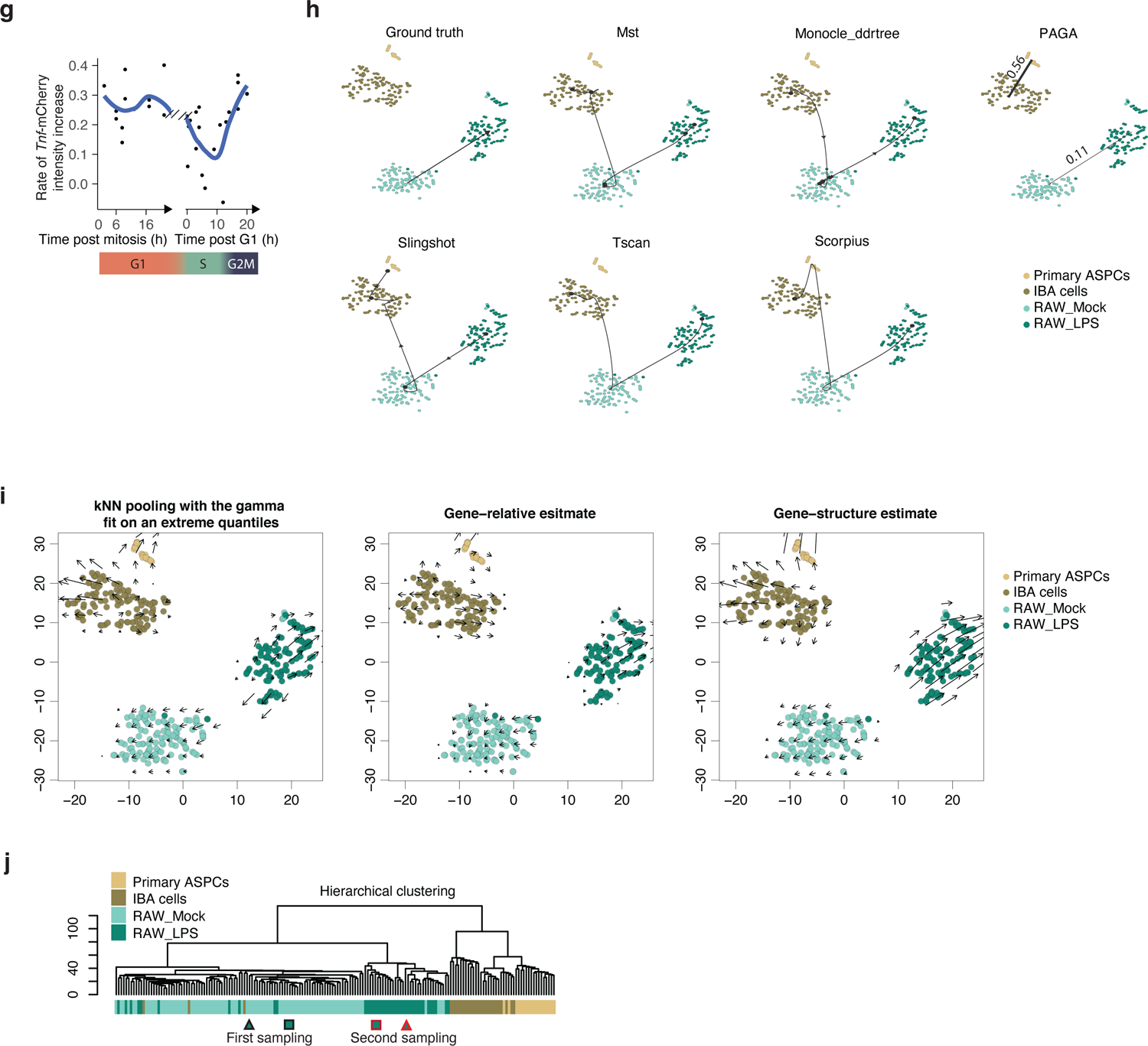
(**a**) Additional results as those shown in Figure 4c, showing a linear relationship between the time post-LPS treatment (within a window of 3 to 7.5 hours post-LPS treatment) and the *Tnf*-mCherry fluorescence intensity (log transformed). (**b**) A linear regression model was used to predict the intercept (basal *Tnf*-mCherry intensity) calculated from data shown in Figure 4c and **Supplementary 4a** based on ground-state gene expression data recorded by Live-seq. The most variable genes from both Live-seq and scRNA-seq data were used while genes that were not detected in Live-seq samples were removed. The *Tnf* gene is highlighted. (**c**) *Gsn* expression (basal) anti-correlates with the rate of *Tnf*-mCherry fluorescence intensity increase. R^2^ and Pearson’s r values are listed. (**d**) *Nfkbia* is one of the most variably expressed LPS-NF-kB pathway genes in multiple primary macrophage cell populations. The expression and the variance of all expressed genes are shown, with genes of the KEGG NF-κB signaling pathway that operate downstream of the TLR4 receptor highlighted. (**e**) The S phase score anti-correlates with *Tnf* expression in conventional scRNA-seq data. (**f**) Cell cycle phases of individual cells were determined using the Fucci reporter miRFP-hCdt1. The Fucci reporter and LPS-induced *Tnf*-mCherry intensities of each individual cell are shown separately, while all cells assigned to a specific cell cycle phase are merged into one meta-plot (at the top of each category) to facilitate direct comparisons. The time of LPS treatment is defined here as 0. Black dots indicate the mitosis time point. The curves of Fucci reporter after LPS treatment are not considered and are therefore shown in a lighter color. The G1/S boundary is inferred from the time point at which the Fucci reporter intensity drops. Cells that underwent mitosis but did not yet reach the G1/S boundary were annotated as G1 cells. Given the lack of a clearly discernable S/G2M boundary, cells were assigned to either the S or G2M phase based on the post G1/S boundary timing. (**g**) The rate of *Tnf*-mCherry fluorescence intensity increase (slope) between 3 to 7.5 hours post-LPS treatment was calculated based on profiles in Figure 4h. The Fucci reporter was used to time cells based on the G1/S boundary. However, mitosis was used to specifically time G1 cells, as the latter did not yet reach the G1/S boundary (which can be detected using the Fucci reporter), rendering them more difficult to annotate. (**h**) Trajectory predictions based on conventional scRNA-seq data using distinct approaches with default settings as contained in the dynverse package. The values in the PAGA plot each represent the probability that the respective clusters are connected. (**i**) Trajectory prediction using the RNA velocity approach. Different strategies including “kNN pooling with gamma fit on extreme quantiles”, “Gene-relative estimate”, and “Gene-structure estimate” were tested. (**j**) The same as shown in **Supplementary** Figure 2g, but with the two sequentially sampled cells highlighted using respectively a rectangle and triangle. A black shape outline represents unstimulated cells, whereas a red shape outline represents LPS-stimulated cells.

**Supplementary Table 1.** Differentially expressed genes in Live-seq samples.

**Supplementary Table 2.** Differentially expressed genes in scRNA-seq samples.

**Supplementary Table 3.** Linear regression model predicting the basal *Tnf-*mCherry intensity using Live-seq data.

**Supplementary Table 4.** Linear regression model predicting the rate in *Tnf*-mCherry fluorescence intensity increase using Live-seq data.

**Supplementary Table 5.** Genes found in negative controls.

**Supplementary Text** is available for this paper.

