## Supplementary Text 1 for "Genome-wide molecular recording using Live-seq"

Since Smart-seq2 is the most sensitive RNA-seq method to detect low input RNA (Ziegenhain et al., 2017), we tested whether it could amplify cDNA at this picogram scale. While successful in ampfying cDNA when the input total RNA was above 5 pg, it failed when input RNA was 1 pg (**Supplementary Figure 1a**). However, 10% of the amount of cDNA reversed transcripted from 10 pg of input RNA (equivalent to 1 pg input RNA) could be amplified (**Supplementary Figure 1a**), suggesting that the reverse transcription product from 10 pg input RNA is more than 10 times than that from 1 pg input RNA. We thus reasoned that the reverse transcription step, rather than the PCR step, is the main reason for the failure. We thus focused on the optimization of the reverse transcription process. We first compared different reverse transcriptase. Among the 5 enzymes tested, only Maxima H Minus Reverse Transcriptase produced significant output with 1 pg total RNA, but showed a similar background signal compared to the negative control (0 pg RNA) (**Supplementary Figure 1b**). Size analysis using a fragment analyzer revealed a hedgehog like profile of both types of cDNA, indicating an amplification of adaptor concatemers (**Supplementary Figure 1c**). However, cDNA from 1 pg RNA showed additional peaks at around 1.1 and 1.8kb (**Supplementary Figure 1c**). We hypothesized that 1 pg RNA could be efficiently reverse transcripted in this condition but this process is hindered by the large amount of adaptor concatemers. We thus tried to reduce the adaptor concatemer by modifying the template-switching oligonucleotide (TSO) (Kapteyn et al., 2010) . Modification of the 5’ end of the TSO by introducing a hairpin (hairpin-TSO) or non-natural nucleotides (iso-TSO) (Kapteyn et al., 2010) did not generate concatemers but also did not yield cDNA (**Supplementary Figure 1d, Methods**), while biotin modified TSO (biotin-TSO) (Islam et al., 2012) largely reduced the concatemer background and did not compromise the cDNA yield. We then tested combinations of different amounts of the TSO, oligo-dT and the reverse transcriptase, to further reduce the background and enhance net cDNA yield (**Supplementary Figure 1e**). The condition with 0.8 µM biotin-TSO, 1 µM oligo-dT and 0.1 µl Maxima H Minus Reverse Transcriptase (200 U/µl) outperformed (**Supplementary Figure 1e-g**), and was thus defined as "the modified Smart-seq2 workflow" from hereon in the main manuscript. We then sequenced the libraries derived from modified Smart-seq2-generated cDNA from 1 pg and 0 pg (negative control) of total RNA. For 1 pg of input RNA, the uniquely mapped rate (rate of the read mapped to the genome among total reads) and exon mapped rate (rate of reads mapped to exon among unique mapped reads) were more than 0.6, with more than 1300 genes detected, while the 0 pg RNA library showed a low uniquely mapped rate, exon mapped rate and small amount of detected genes (**Supplementary Figure 1h**). The sequences derived from oligo-dT and TSO were overrepresented in the 0 pg RNA library (**Supplementary Figure 1i-k**). The top 20 genes absorbed most of the mapped reads of these libraries (**Supplementary Figure 1k**). These are likely due to sequencing errors or mis-mapping of the A/T rich region and thus were not included in downstream data analyses.

Islam, S., Kjällquist, U., Moliner, A., Zajac, P., Fan, J.-B.B., Lönnerberg, P., and Linnarsson, S. (2012). Highly multiplexed and strand-specific single-cell RNA 5’ end sequencing. Nat Protoc *7*, 813–828.

Kapteyn, J., He, R., McDowell, E.T., and Gang, D.R. (2010). Incorporation of non-natural nucleotides into template-switching oligonucleotides reduces background and improves cDNA synthesis from very small RNA samples. BMC Genomics *11*, 413.

Ziegenhain, C., Vieth, B., Parekh, S., Reinius, B., Guillaumet-Adkins, A., Smets, M., Leonhardt, H., Heyn, H., Hellmann, I., and Enard, W. (2017). Comparative Analysis of Single-Cell RNA Sequencing Methods. Molecular Cell *65*, 631-643.e4.
